# Identification of decondensed large-scale chromatin regions by TSA-seq and their localization to a subset of chromatin domain boundaries

**DOI:** 10.1101/2021.04.06.438686

**Authors:** Omid Gholamalamdari, Liguo Zhang, Yu Chen, Andrew Belmont

## Abstract

Large-scale chromatin compaction is nonuniform across the human genome and correlates with gene expression and genome organization. Current methodologies for assessing large-scale chromatin compaction are indirect and largely based on assays that probe lower levels of chromatin organization, primarily at the level of the nucleosome and/or the local compaction of nearby nucleosomes. These assays assume a one-to-one correlation between local nucleosomal compaction and large-scale compaction of chromosomes that may not exist. Here we describe a method to identify interphase chromosome regions with relatively high levels of large-scale chromatin decondensation using TSA-seq, which produces a signal proportional to microscopic-scale distances relative to a defined nuclear compartment. TSA-seq scores that change rapidly as a function of genomic distance, detected by their higher slope values, identify decondensed large-scale chromatin domains (DLCDs), as then validated by 3D DNA-FISH. DLCDs map near a subset of chromatin domain boundaries, defined by Hi-C, which separate active and repressed chromatin domains and correspond to compartment, subcompartment, and some TAD boundaries. Most DLCDs can also be detected by high slopes of their Hi-C compartment score. In addition to local enrichment in cohesin (RAD21, SMC3) and CTCF, DLCDs show the highest local enrichment to super-enhancers, but are also locally enriched in transcription factors, histone-modifying complexes, chromatin mark readers, and chromatin remodeling complexes. The localization of these DLCDs to a subset of Hi-C chromatin domain boundaries that separate active versus inactive chromatin regions, as measured by two orthogonal genomic methods, suggests a distinct role for DLCDs in genome organization.

## Introduction

With two meters of DNA inside every human cell, a principal question in genome biology is how cells pack this material inside the ∼10 µm diameter nucleus. Based on extensive previous studies, we know that isolated chromatin folds under certain salt conditions, including physiological univalent cation concentrations, into ∼20-40 nm diameter fibers, termed “the 30 nm” higher-order chromatin fiber (Bian and Belmont 2012; Grigoryev and Woodcock 2012).

Physiological concentrations of divalent and multivalent cations, however, lead to an aggregation/oligomerization of isolated or reconstituted chromatin in vitro, dependent on specific histone tail modifications as well as distance-dependent effects induced by local binding of transcriptional coactivators such as p300, as measured by analytical ultracentrifugation (Schwartz et al. 1996; Shogren-Knaak et al. 2006; Lu et al. 2006; McBryant et al. 2009; Szerlong and Hansen 2011; Watanabe and Peterson 2010). More recently, this condensation of chromatin oligonucleosomes has been likened to a liquid-liquid phase separation of 10 nm chromatin fibers (Maeshima et al. 2016; Gibson et al. 2019).

Using both classical heavy-metal as well as several DNA-specific stains, transmission electron microscopy (TEM) of chromosomes and nuclei reveals most of the genome is folded within condensed chromatin regions much larger than 30nm in typical mammalian somatic cells, as reviewed previously (Belmont 2014). For reasons that remain unclear, a few more recent studies have visualized dispersed chromatin, without evident organization into large-scale chromatin domains (Ricci et al. 2015; Ou et al. 2017). However, both direct visualization of chromatin in live-cells (Robinett et al. 1996; Li et al. 1998; Tumbar et al. 1999; Hu et al. 2009; Deng et al. 2016), including super-resolution imaging approaches (Nozaki et al. 2017), and TEM on samples fixed using high-pressure freezing (Hoffman et al. 2020) yield results consistent with the previous observations of large-scale chromatin domains. Interestingly, the increased compaction of large-scale chromatin domains with increasing divalent and polycation concentrations observed in both mitotic and interphase chromosomes in situ parallels the increased aggregation/oligomerization observed for oligonucleosomes in vitro. Moreover, increased large-scale chromatin compaction is observed in live cells exposed to hypertonic media (Albiez et al. 2006), while increased chromatin compaction during mitosis has recently been linked to physiological increases in free Mg++ cellular concentration (Maeshima et al. 2018). Cell cycle progression is accompanied by progressive apparent unfolding and then refolding of these large-scale chromatin domains during progression through interphase and then back into mitosis (Belmont and Bruce 1994; Kireeva et al. 2004).

Within individual nuclei and chromosomes, variations in large-scale chromatin compaction appear in different chromosome regions tied to changes in gene activation. Targeting transcriptional activator domains to a gene-amplified, heterochromatic domain led to a striking large-scale chromatin decondensation through an apparent uncoiling of a large-scale chromatin fiber (Tumbar et al. 1996, 1999; Carpenter et al. 2005). Using stably integrated, BAC transgene arrays, which reconstitute per copy gene expression levels within several-fold of their endogenous gene counterparts, a 2-3 fold large-scale chromatin decondensation was observed after transcriptional induction of several inducible gene loci (Hu et al. 2009). In all cases, average linear compaction ratios remained ∼10-fold or more, higher than predicted for 30-nm chromatin fibers even after transcriptional induction, even as observed directly in living cells.

Examination of endogenous chromosome regions using FISH, although subject to some decompaction of chromatin structure as a result of DNA denaturation (Solovei et al. 2002; Hepperger et al. 2007), similarly showed decondensation of endogenous gene loci following transcriptional induction, while maintaining an overall level of compaction significantly higher than expected for 30-nm fibers (Volpi et al. 2000; Mahy et al. 2002; Müller et al. 2004; Christova et al. 2007). Thus, transcription for typical genes in many mammalian cell types occurs on a condensed template. Similarly, pulse-chase labeling experiments have revealed that DNA replication takes place on the exterior surface of a condensed large-scale chromatin template (Deng et al. 2016).

Despite several existing genome-wide technologies to assess chromatin compaction, none directly measure this highest level of large-scale chromatin compaction. Several methods-DNase-Seq, MNase-Seq, ATAC-Seq-measure the accessibility of DNA to modifying enzymes (Boyle et al. 2008; Giresi et al. 2007; Buenrostro et al. 2013). Other strategies rely on the physical or chemical properties of chromatin and its response to crosslinking agents (e.g., FAIRE-seq) or gradient centrifugation (Gilbert et al. 2004). These approaches have two main problems. First, they measure chromatin compaction indirectly and second, they measure compaction at lower levels of folding, which may not be equivalent to what we see under the microscope. Microscopic experiments, on the other hand, define chromatin compaction as the DNA content per unit volume or as a linear compaction ratio relative to B-form DNA as observed on intact chromosomes in nuclei.

To reconcile microscopic versus genome-wide measurement of compaction, we adapted a recently developed methodology, TSA-seq. This method estimates distances of chromosome loci relative to a nuclear compartment such as nuclear speckles (Chen et al. 2018; Zhang et al. 2020). Here we use the first-order derivative of the TSA-seq signal, proportional to both the local chromatin compaction and the chromosome orientation relative to the TSA staining target, to identify “high-slope” genomic regions with unusually decondensed large-scale chromatin structure, which we have termed <u>D</u>econdensed <u>L</u>arge-scale <u>C</u>hromatin <u>D</u>omains, or “DLCDs”.

DLCDs map adjacent to Hi-C-defined chromatin domain boundaries and are enriched for active chromatin modifications, transcription factors, and structural components of chromatin.

## Results

### How TSA-seq estimates chromatin compaction based on relative cytological distances

We hypothesized that the radial distance function of chromosome regions from a reference point inside the cell nucleus (e.g. nuclear speckles) contains chromatin compaction information, whose first derivative should be proportional to the local chromatin compaction and chromosome orientation relative to the reference nuclear compartment (assuming a random orientation of chromatin with respect to the reference point). Therefore, a decondensed 200 kb region travels a greater distance from nuclear speckles (Fig. 1A, right, orange) compared to a more condensed region of similar DNA length (Fig. 1A, right, green), and should have a higher slope in the distance function (Fig. 1A, left).

**Figure 1.**
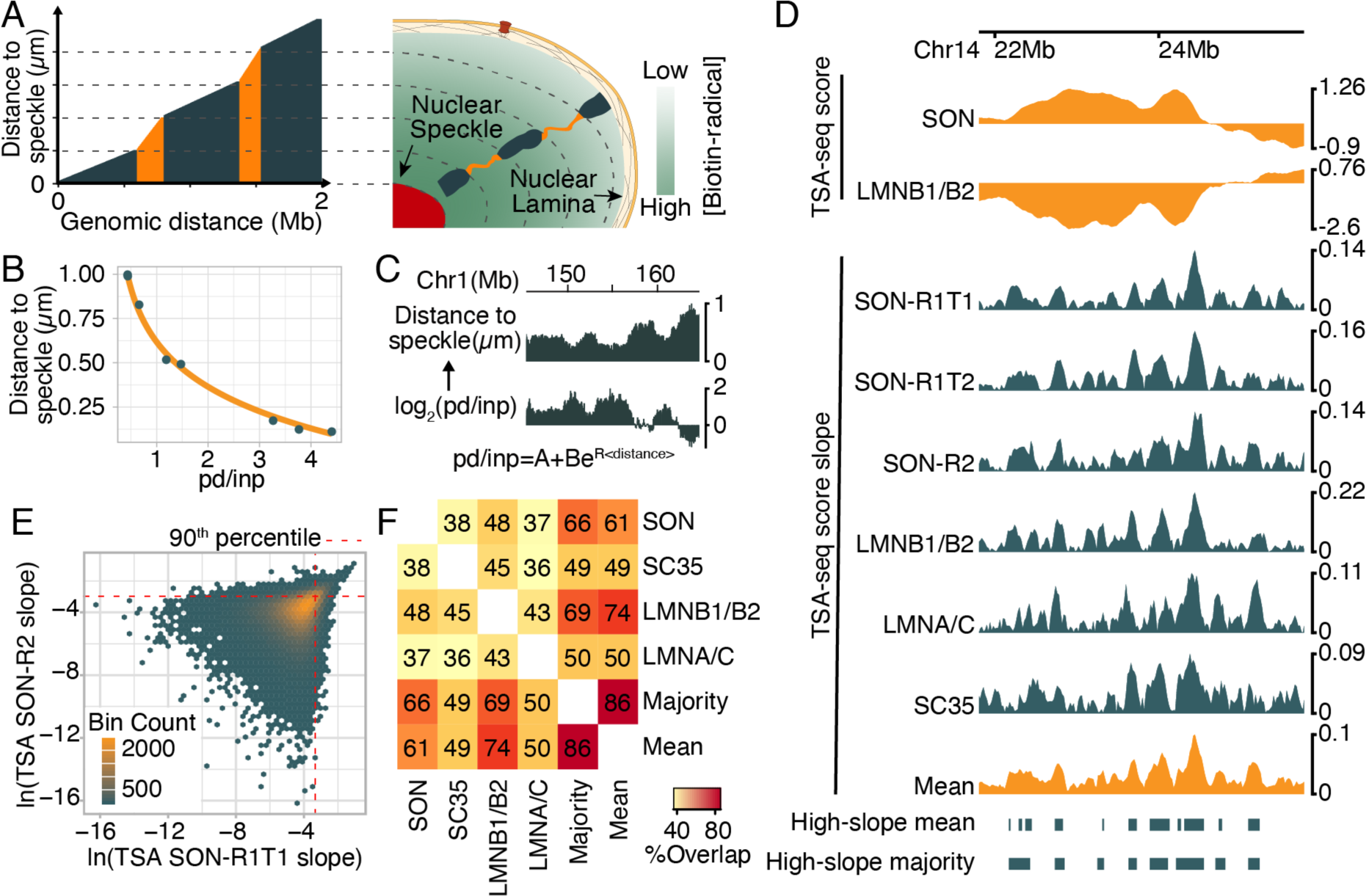
Adapting TSA-seq to measure decondensed large-scale chromatin domains. **(A)** Distance from nuclear speckle plotted for a hypothetical genomic region (left) and schematic representation of expected large-scale chromatin domains (right). Decondensed Large-scale Chromatin Domains (DLCDs) highlighted in orange have higher slopes compared to green genomic regions, implying a relative decondensation of orange compared to green chromosome domains. **(B)** Exponential fitting of 3D-FISH distance from nuclear speckles versus TSA-seq pd/inp (K562 FISH data is reproduced from Chen et al). **(C)** Speckle distances (chr1:145-165 Mbp) for K562 calculated from the inverse of the exponential fit. **(D)** TSA-seq score and slope genomic plots in K562 cells for a region of Chr14. Top to bottom tracks: nuclear speckle (SON) and nuclear lamina (LMNB1/B2) TSA-seq scores, absolute values of TSA-seq score slopes (absolute magnitude) for SON TSA-seq replicate 1 technical replicate 1 (R1T1), SON replicate 1 technical replicate 2 (R1T2), SON replicate 2, LMNB1/B2 replicate 1, nuclear lamina LMNA/C replicate 2, nuclear speckle SC35 replicate 2, average of TSA-seq slopes calculated across all available TSA-seq slope datasets (Mean). Bottom two tracks show high-slope domains identified based either on top decile of the mean of all TSA-seq score slopes (2^nd^ from bottom track) or top decile domains present in the majority of individual datasets (high-slope majority) (bottom track). **(E)** TSA-seq score slopes are reproducible for high slope values. Binned scatter plot showing the number of bins with given values of the natural log of SON TSA-seq score slope (absolute value of derivative) calculated from replicate 1 (rep1) (x-axis) versus replicate 2 (rep2) (y-axis) TSA-seq datasets. The red dashed lines show the 90^th^ percentile of the derivative values in each replicate. **(F)** Percent overlap (Jaccard index) between high-slope domains identified based on individual TSA-seq slope datasets and merged high-slope domains (mean and majority).

TSA-seq estimates this distance function by exploiting an established cytological method, Tyramide Signal Amplification, which was designed to amplify the signal from antibody staining approaches (Bobrow et al. 1989). Fixed cells are stained with a primary antibody against a protein enriched in the target compartment (e.g., SON or SC-35 for nuclear speckles). The primary antibody is stained with a secondary antibody conjugated to horseradish peroxidase (HRP), which catalyzes generation of tyramide-biotin free radicals. These free radicals diffuse from their secondary antibody source, creating a steady-state concentration gradient of tryamide free radicals that decays exponentially with distance from the HRP source. Tyramide-biotin free radicals directly label DNA, thus creating a distance-dependent biotinylation of DNA proportional to the local concentration of tyramide free-radicals. Upon purification of biotin-labeled DNA, sequencing, and alignment to the genome, one can compute the pull-down (pd) over input (inp) ratio, pd/inp, for all genomic regions. 3D fluorescent in-situ hybridization (3D-FISH) confirms that the pd/inp ratio has an exponential relationship with mean 6 distance to nuclear speckles (Fig. 1B) (FISH data is reproduced from (Chen et al. 2018); Supplemental Table S1). Using this exponential relationship, we can convert the TSA-seq pd/inp ratios into estimated cytological distances from a nuclear compartment such as nuclear speckles, which are approximated as point-source generators of tyramide free radicals (Fig. 1C).

**Table 1.** Summary of 3D-FISH experiments

| Test Probe | Validation cell-line | Slope percentile<br>(median, K562) | Test probe Feret<br>(# of loci) | Shared probe<br>Feret (# of loci) |
| --- | --- | --- | --- | --- |
| RP11-49O17 | K562 | >99 | 0.54µm (381) | 0.49µm (340) |
| RP11-33O9 | K562 | >99 | 0.56µm (416) | 0.49µm (748) |
| RP11-8P9 | K562 | 82 | 0.51µm (309) | 0.45µm (358) |
| RP11-1008C9 | K562 | 38 | 0.47µm (383) | 0.45µm (591) |
| RP11-33O9 | HCT116 | >99 | 0.48µm (365) | 0.43µm (306) |
| RP11-173P16 | HCT116 | 48 | 0.58µm (408) | 0.58µm (459) |
| CTD-2503D10 | HCT116 | 52 | 0.48µm (433) | 0.45µm (319) |

Here we use the first derivative of the normalized TSA-seq score, that involves the natural logarithm of the pd/inp ratio. This TSA-seq score should be approximately a linear function of distance from the source, given the exponential dependence of distance on the pd/inp ratio. Therefore, similar to the distance function, its first derivative should report on chromatin compaction and orientation relative to the reference point.

### High TSA-seq derivative is reproducible

To numerically calculate the TSA-seq slope, we opted to smooth the published TSA-seq datasets from K562 cells (Chen et al. 2018). The TSA-seq normalized scores were calculated for 20 kb bins across the genome. Data values over a sliding window of different sizes (5 to 41 bins) were fitted to cubic polynomials, and the central bin value replaced by the calculated value of this polynomial. As expected, using different numbers of genomic bins for fitting results in different degrees of smoothing (Supplemental Fig. S1). The optimal window size for smoothing was guided by comparing reproducible features of SON TSA-seq replicates. We chose a window size of 420 kb (21 bins of 20kb) for further analysis. Subsequently, the TSA-seq derivative of the central genomic bin was calculated as the derivative of the cubic polynomial fitted to the original TSA-seq score data. Because the derivative sign denotes only the chromatin trajectory direction relative to the viewpoint (i.e., TSA-seq staining target), we used the absolute magnitude of the derivative (hereafter called slope) as a metric proportional to large-scale chromatin compaction and orientation. We calculated this slope for all available TSA-seq datasets in K562, including SON and SC35 (nuclear speckles) and LMNB1/B2 and LMNA/C (nuclear lamina) (Fig. 1D).

Comparing SON TSA-seq replicates, we observed increased reproducibility for higher values of the TSA-seq score slope, particularly the highest 10% of slope magnitudes genome-wide (Fig. 1E and Supplemental Fig. S2). We therefore focused our analysis on these top 10% of TSA-seq slope values, hereafter defining them as “high-slope” regions.

Using this 10^th^ percentile threshold, we identified 4116 high-slope regions in SON TSA-seq replicate 1 and 4508 high-slope regions in SON TSA-seq replicate 2, with 2174 of these regions identified in both replicates (Supplemental Fig. S3). Mean and median sizes of these high-slope regions are 69 and 60 kbp, respectively. Of the 4253 high-slope regions identified in only one SON TSA-seq replicate, 1106 were detected as high-slope using a top 15^th^ percentile slope threshold. Similarly, using SC35 TSA-seq, we obtained a similar set of 4895 high-slope regions (Fig. 1F and Supplemental Fig. S3), with 1725 overlapping with replicate 1 SON TSA-seq high-slope regions.

High-slope regions are robust to the choice of the TSA-seq target nuclear structure as they also can be identified using nuclear lamina TSA-seq datasets (Fig. 1D, 1F, and Supplemental Fig. S2-3), with an ∼50% overlap between high-slope regions detected using SON versus LMNB1/B2 TSA-seq datasets (Fig. 1F and Supplemental Fig. S3). Lowering the threshold to 85% for the percentile slope cutoff used in the lamin B1/B2 TSA-seq data set increases this overlap to 60%.

To consolidate high-slope regions detected by various TSA-seq datasets and to minimize false positive rates in detecting DLCDs, we merged high-slope regions using two different strategies. First, we calculated the average derivative of available TSA-seq slope datasets in K562 and called high-slope regions using a 10% cut-off value in the genomic percentile for these averaged values (Fig. 1D and 1F; high-slope mean). Second, we used a “consensus counting” approach: high-slope regions were called for each dataset separately using a 10% threshold in TSA-seq derivative genomic percentile. A region was then classified as high-slope if it was above this 10% threshold in a simple majority of all the TSA-seq datasets (Fig. 1D-F; high-slope majority). There is an 86% overlap in the high-slope regions identified by these two strategies; additionally, high-slope regions identified by either of these two strategies overlap >50% with high-slope regions identified in each of the individual TSA-seq datasets (Fig. 1F and Supplemental Fig. S3). We used the high-slope mean dataset for further genomic data analysis (See Supplemental Table S2 for the genomic coordinates of these regions).

### TSA-seq high-slope regions identify decondensed large-scale chromatin domains

To test whether TSA-seq high-slope regions correspond to genomic regions with decondensed large-scale chromatin structure, we used 3D-FISH to compare high-slope genomic regions with control regions in K562 cells.

Due to the required melting of DNA secondary structure used in most FISH protocols, as well as an additional deproteinization step in the 3D-FISH protocol, typically the chromosome structure detected after FISH is noticeably decondensed, as visualized by wide-field and electron microscopy (Robinett et al. 1996; Solovei et al. 2002; Hepperger et al. 2007).

Additionally, we observed that longer storage of coverslips in formamide-containing buffers at +4°C, which is typically done to increase hybridization efficiency, appeared to result in an increased projected area of the FISH signal. Therefore, we included an internal control BAC (RP11-1047B3) probe labeled in a separate imaging channel to correct for chromatin compaction variations induced by uncontrolled variability in the FISH procedure. This shared BAC probe (Supplemental Fig. S4C) corresponds to a lamina-associated domain (LAD).

We selected 6 test BACs-3 mapped to genomic regions with high (top 10 percentile) and 3 with lower levels of TSA-seq slope (38-52 percentile)-to compare against the internal standard and each other (Table 1 and Supplemental Fig. S4A-B). BAC test probes were labeled with DIG-dUTP and the internal control BAC labeled with biotin-dATP. FISH signals corresponding to high-slope regions were significantly more elongated compared to the FISH signals corresponding to both the internal control BAC and the BACs from “low-slope” regions (Fig. 2A).

**Figure 2.**
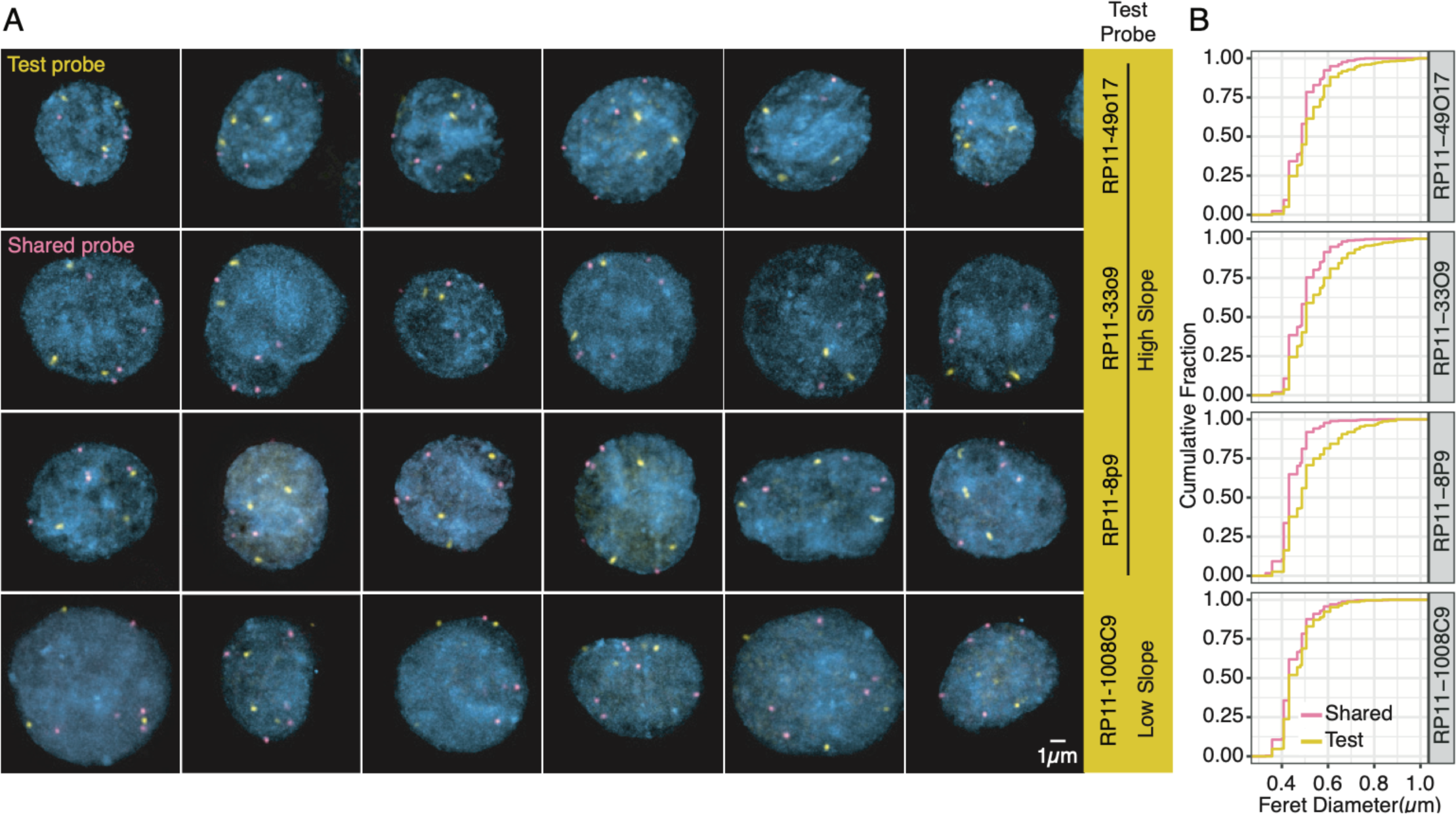
High-slope domains are decondensed relative to low-slope control regions. **(A)** 3D-FISH in K562 cells shows high-slope regions appearing as larger, more elongated signals in ∼50% of cells compared to internal standard and low-slope region. Test BAC probes (yellow) are from high-slope regions (top 3 rows) or from a low-slope region (bottom row). Shared BAC probe (magenta) corresponding to LAD region is used as an internal standard to control for varying preservation of chromosome structure after FISH procedure. Nuclei are counterstained with DAPI (cyan). Images are maximum intensity projections of entire nuclei. Scale bar = 1µm. **(B)** Cumulative distribution of 3D-FISH signal Feret diameters (longest line through FISH signal). Fraction of alleles (y-axis) are shifted towards larger diameters (x-axis) for test FISH probes (yellow) compared to shared probe (magenta line) for high-slope probe regions (top 3 panels), whereas test probe from low-slope region shows same distribution as internal standard (bottom panel). n≥200 (350) K562 cells (alleles) were imaged for each test probe.

To better compare these FISH signals, we measured their Feret diameter (longest line segment intersecting the segmented image convex hull) (Table 1). Greater than 100 nuclei with greater than 300 alleles were measured for each BAC test probe. Background noise was subtracted and then each FISH signal was segmented using a local threshold equal to 50% of its peak intensity. Feret diameter distributions for both high and low slope BAC probe FISH signals were similar to the internal standard BAC FISH signals over the lower two quantiles of the distribution (Fig. 2B, Supplemental Fig. S5). However, the upper two quantiles of the distributions for the high-slope regions displayed significantly more decondensed FISH signals relative to the internal standard (Figs. 2B, Supplemental Fig. S5). In contrast, Feret diameter distributions for all low-slope FISH signals were similar across all four quantiles to the internal standard (Fig. 2B, Supplemental Fig. S5). We performed Welch two sample t-test between the shared and test probes’ Feret diameter. The differences between the Feret diameter means for the high-slope and shared probes (≥0.05 µm, p < 0.0001) and the low-slope shared probes (∼0.02 µm, p<0.0001) were both significant, although the mean difference was very small between the shared and low-slope probes.

To rule out the possibility that the observed Feret diameter differences between high-slope versus internal standard and low slope FISH signals are due to earlier DNA replication of the high-slope versus low slope genomic regions, which introduces a bias towards larger FISH signals due to an increased percentage of replicated alleles in the cell population, we performed FISH in HCT116 cells pulsed briefly with EdU, a thymidine analog. We compared Ferret diameters of FISH signals from EdU positive, S-phase cells versus EdU-negative, G1 and G2 cells, which are mostly G1 cells (70% G1 versus 5% G2+M as measured by flow cytometry). In both the S-phase and G1/G2 populations we observed a similar increase in Feret diameters for the top two quartiles of high-slope BAC FISH signals, similar to the distributions of the entire cell population, whereas Feret diameters for low-slope BAC FISH signals were similar to the internal standard across the entire percentile range (Supplemental Fig. S5).

In summary, these high-slope regions correspond to genomic regions showing increased large-scale chromatin decondensation in ∼50% of the allelic population, as visualized by 3D-FISH wide-field microscopy. We therefore refer to these high-slope regions as <u>D</u>econdensed <u>La</u>rge-scale <u>C</u>hromatin <u>D</u>omains, or DLCDs.

### DLCDs map to boundaries of chromatin topological domains

Examination of the chromosomal distributions of TSA-seq high-slope regions revealed that they map close to the boundaries of the triangular domains identified by Hi-C contact maps (Fig. 3A and 4A). Hi-C detects two main compartments, termed A and B, in the genome (Lieberman-Aiden et al. 2009). Compartment A, enriched for transcriptionally active genes, and compartment B, enriched for transcriptionally inactive genes and gene deserts, correspond closely with the iLADs and LADs, respectively, identified by DamID (van Steensel and Belmont 2017). Hi-C topological maps have been further partitioned into sub-compartments, Topological Associated Domains (TADs), and looped-domains, with and without CTCF-anchors (Nora et al. 2012; Dixon et al. 2012; Rao et al. 2014; Fernandez et al. 2020).

**Figure 3.**
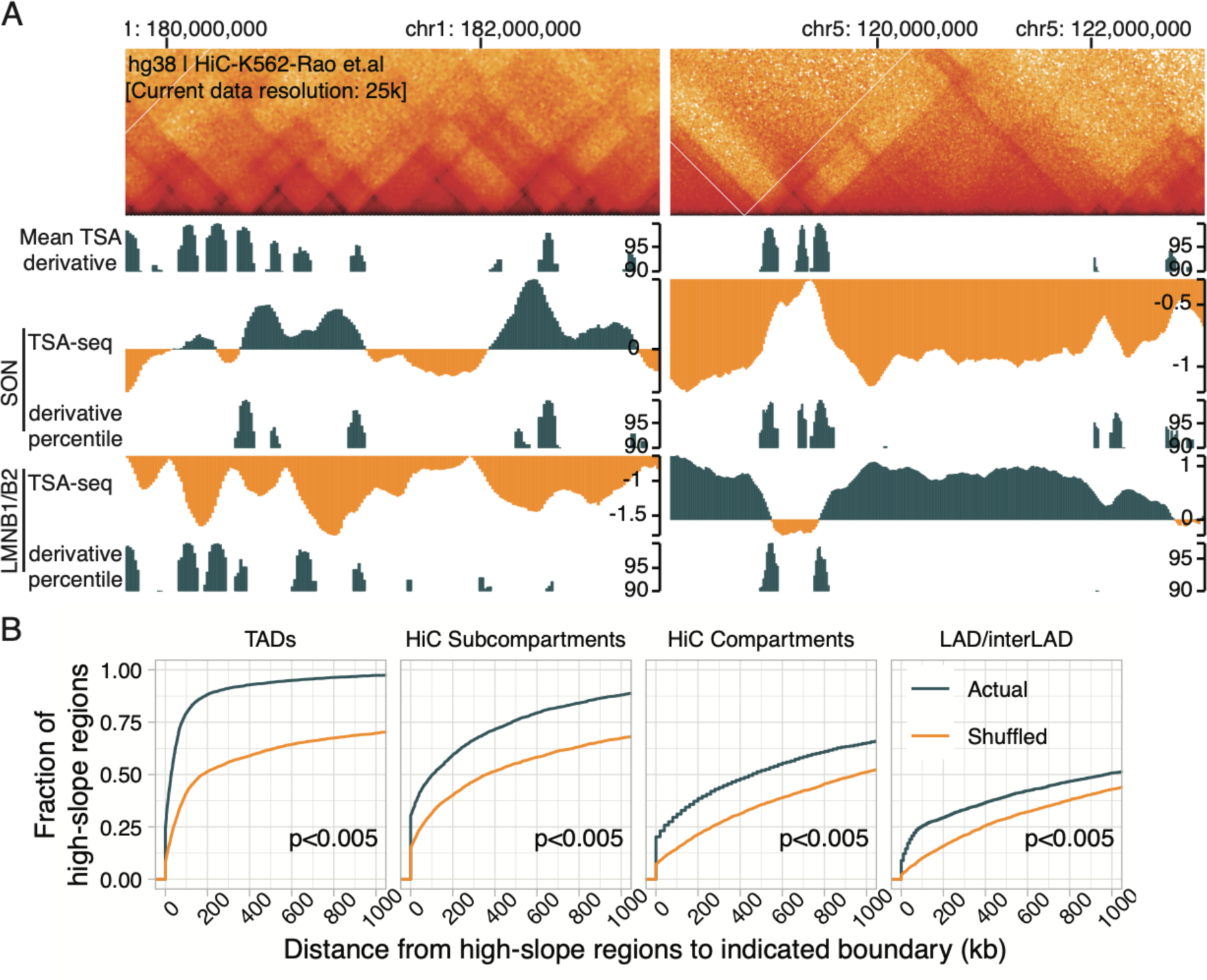
TSA-seq high-slope regions map close to chromatin domain boundaries. (A) Top to bottom: High-slope regions in K562 cells map to the corners of a subset of the “triangular” chromosome domains identified by Hi-C (top panels, 5 kb Hi-C resolution); Top decile of mean of TSA-seq slopes calculated across multiple TSA-seq datasets; SON TSA-seq score (top, green and orange) versus top decile of slope (green); LMNB1/B2 TSA-seq score (top, green and orange) versus top decile of slope (green). (B) Cumulative fraction (y-axis) of the high-slope regions (green line) versus shuffled domains (orange line) as function of distance (x axis, kb) to the boundaries in K562 cells of (left to right): Hi-C TADs, subcompartments, and compartments as well as lamin B1 DamID LADs. P-values were measured empirically by shuffling high-slope regions across the genome 200 times.

**Figure 4.**
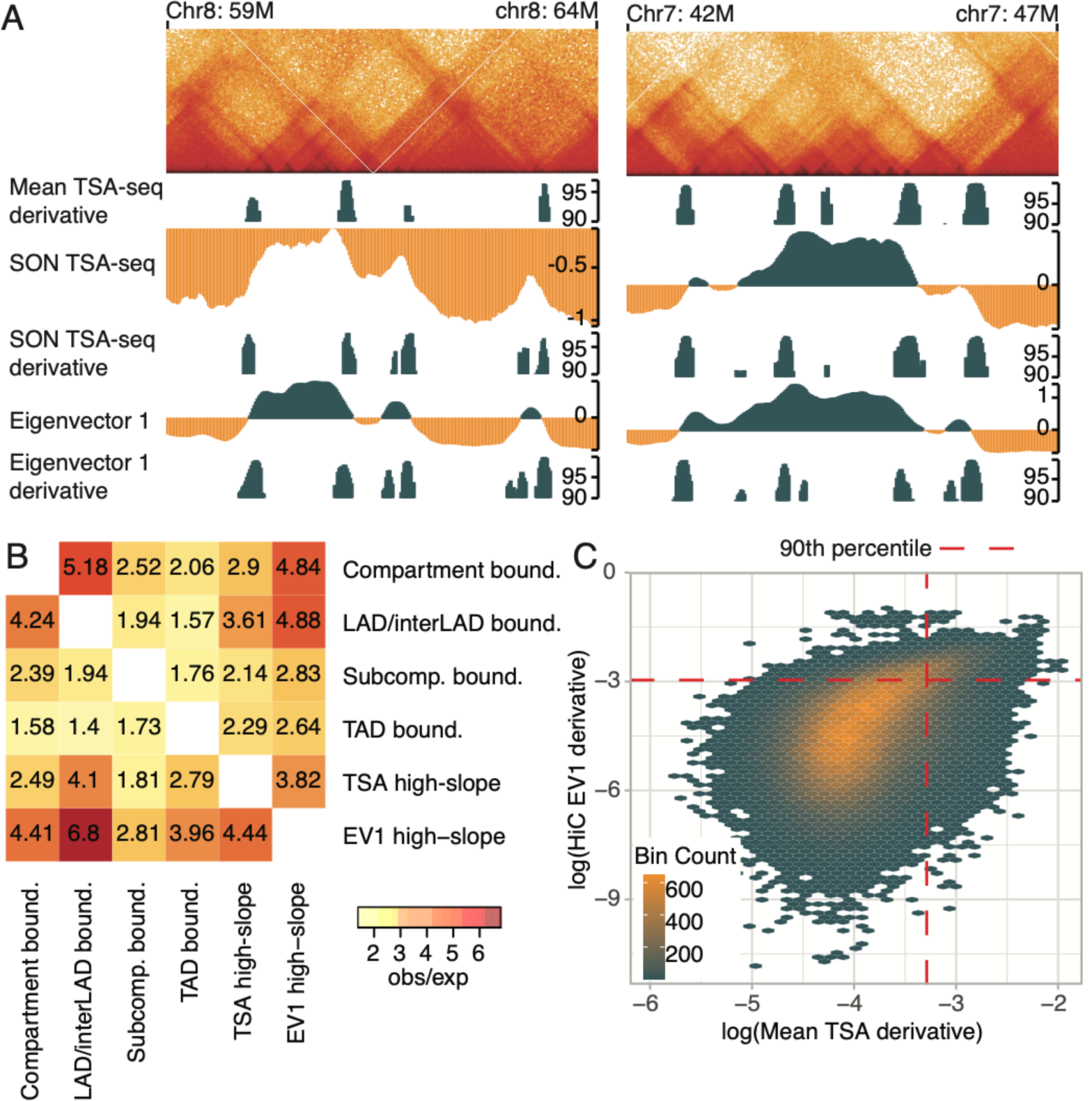
SON TSA-seq and Hi-C EV1 high-slope regions are highly correlated. **(A)** Top to bottom for Chr8 region (left) and Chr7 region (right): Hi-C (K562) interaction map (5kb resolution); Top decile of mean of TSA-seq slopes calculated across multiple TSA-seq datasets (Mean TSA-seq slope); SON TSA-seq score; Top decile shown of SON TSA-seq score slope; Hi-C eigenvector 1 (EV1); Top decile of Hi-C eigenvector 1 slope. **(B)** Observed overlap/expected overlap probability for TSA-seq and EV1 high-slope domains with other chromatin domain boundaries (top to bottom and left to right): Hi-C compartments, LAD/interLAD, Hi-C subcompartments, TAD, TSA-seq high slope, Hi-C EV1 high slope. For the bottom triangle columns are references and the row labels are the queries (i.e., the bottom left number denotes the observed/expected overlap probability of EV1 high-slope domains overlapping with the indicated chromatin domain boundaries) **(C)** Correlation between mean of TSA-seq score slopes (averaged over multiple datasets) and Hi-C EV1 slopes, especially for top decile values of both. Binned scatter plot of the values for TSA-seq (x-axis) and Hi-C EV1 (y-axis) slopes.

To correlate the genomic locations of DLCDs with the topological domains detected by Hi-C, we defined the “center” of high-slope regions as the local maxima of the TSA-seq slope in each region. Greater than 75% of DLCDs map within 100 kbp of TAD boundaries, significantly higher than the 40% of DLCDs that map within 100 kbp of TAD boundaries in the “shuffled” control (Fig. 3B). Similarly, ∼50%, ∼25%, and ∼25% of DLCDs map within 100 kbp of Hi-C subcompartments, A/B compartments, and LAD/inner-LAD boundaries, respectively, as compared to the ∼25%, ∼10%, and ∼10% of DLCDs that map within 100 kbp of these same respective boundaries in the shuffled control (Fig. 3B and Supplemental Fig. S6).

The first eigenvector (EV1) of the Hi-C contact frequency matrix is typically used to identify the Hi-C A and B compartments, with positive EV1 values defining the A compartment, negative EV1 values defining the B compartment, and zero EV1 values defining the A/B compartment boundaries (Lieberman-Aiden et al. 2009; Imakaev et al. 2012). The Hi-C EV1 strongly correlates with the plaid pattern of the Hi-C interaction matrix, GC percent, DNase hypersensitivity, and DNA replication timing (Ryba et al. 2010; Imakaev et al. 2012).

Visual inspection in the genome browser suggested that DLCDs also appeared to align with high-slope regions of the Hi-C EV1 signal. We confirmed this correlation between TSA-seq and EV1 slopes. We calculated the EV1 at 10 kb resolution to capture fine changes in the EV1 signal and estimated its slope in 420kb window. TSA-seq high-slope regions align at high frequency with EV1 high-slope regions, with boundaries of Hi-C “triangular domains” frequently aligning with both TSA-seq and EV1 high-slope regions (Fig. 4A).

More specifically, although a substantial fraction of TSA-seq high-slope regions coincide with EV1 high-slope values, a significant fraction does not. EV1 high-slope regions predominately map to A/B compartment boundaries, but a fraction map inside Hi-C compartments (Fig. 4A; right panel). EV1 and TSA-seq high-slope regions coincide ∼4 times more frequently than expected by chance. They also both coincide at increased frequencies with LAD/interLAD and Hi-C compartment boundaries (Fig. 4B). TSA-seq and EV1 slope values correlate genome-wide (Spearman correlation, R=0.45), but this correlation is enhanced for the highest slope magnitudes (Fig. 4C); similar to the correlation between the TSA-seq slope values (Fig. 1E and Supplemental Fig. S2A).

### DLCDs separate transcriptionally active and inactive chromatin domains

We used available K562 genome-wide datasets (Supplemental Table S3) to systematically identify characteristics of the DLCDs defined by TSA-seq high-slope domains. These datasets included: 1) genic features: transcription start sites (TSSs), house-keeping genes (HK genes), enhancers, and super-enhancers; 2) ENCODE ChIP-seq of histone modifications, transcription-related factors (“TFs”), and chromatin structural proteins; 3) ENCODE RNA-seq and DNase-seq (see Supplemental Table S3 for a complete list). We generated profile plots by averaging dataset signals as a function of distance from each of the TSA-seq high-slope centers (+/-500 kbp).

TSSs, HK-genes, and histone modifications associated with active transcription were all enriched near TSA-seq high-slope centers while inactive chromatin features (e.g., H3K9me3) were instead depleted (Fig. 5A and Supplemental Figs. S7-8). Of all these features, super-enhancers showed the largest relative enrichments near the TSA-seq high-slope centers, with a >300% increase in peak versus flanking region signals (Fig. 5A, Supplemental Fig. S7). Additionally, insulator protein CTCF and cohesin complex subunits, RAD21 and SMC3, were enriched near TSA-seq high-slope centers (Fig. 5A).

**Figure 5.**
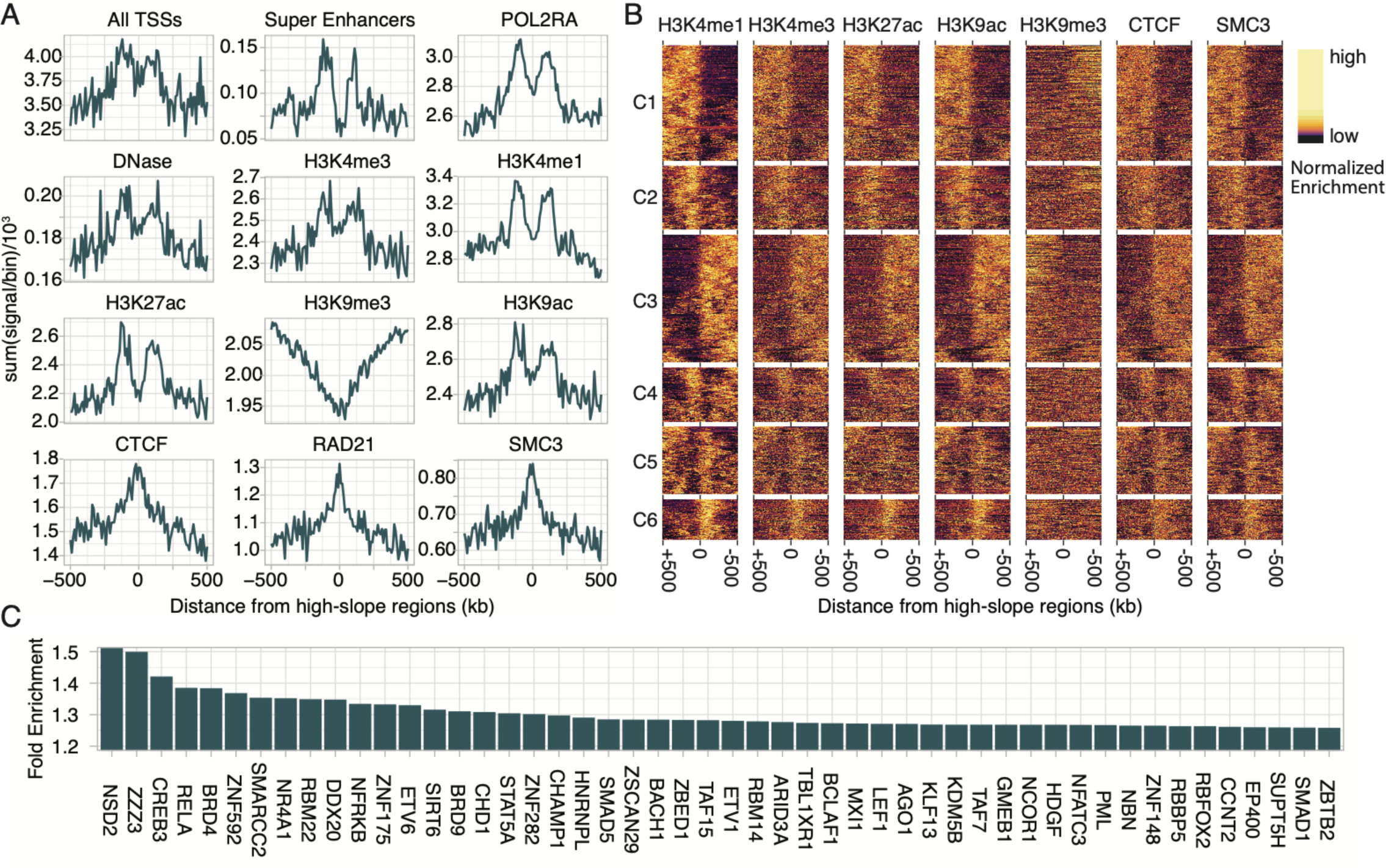
DLCDs are flanked on one side by an enrichment peak for active genomic and chromatin features. **(A)** Averaged profile plots showing genomic features or chromatin ChIP-seq flanking (+/-500 kb) the w), H3K4me3, H3K4me1, H3K27ac, H3K9me3, H3K9ac, CTCF, RAD21, SMC3. Units are feature count per bin for genic features, ChIP-seq peak counts for POLR2A, CTCF, RAD21, and SMC3, and ChIP-seq pulldown/input per bin for various ChIP-seq against histone marks. **(B)** Heatmaps of histone modifications, CTCF, and SMC3 ChIP-seq flanking (+/- 500 kb) the centers of TSA-seq high-slope regions. Heatmap displays pulldown/input ChIP-seq ratios normalized independently for each row to emphasize the local changes of these marks relative to the high-slope centers (0 kb). Heatmap rows are clustered using H3K4me1 data. Data in A and B is binned at 10kb. **(C)** Top 50 ChIP-seq peak enrichment relative to genomic background for regions flanking TSA-seq high-slope domain centers as in (A), calculated for over 300 chromatin related factors (see Supplemental Figure S7 for the original profile plots).

ReMap 2020 (Chèneby et al. 2020) has catalogued 560 ChIP-Seq datasets for 349 unique chromatin associated factors in K562 cells. We found 238 out of these 349 factors with varying levels of enrichment across high-slope regions (Fig. 5A and Supplemental Fig. S8).

Most active chromatin features exhibited a bimodal enrichment distribution, with peaks located symmetrically on each side of the TSA-seq high-slope center. These two peak distributions overlapped partially, creating a dip at the high-slope center, with a peak-to-peak separation of ∼250kb. Exceptions to this rule were CTCF, RAD21, SMC3, and CBX8 whose distributions instead showed a broader unimodal distribution centered over the TSA-seq high-slope regions (Fig. 5A and Supplemental Fig. S7-8). A wide but unimodal overall distribution of CTCF, RAD21, and SMC3 relative to TAD boundaries was reported previously (Dixon et al. 2012).

These bimodal distributions suggest two alternative explanations. 1) Actual enrichment of active chromatin features in regions flanking both sides of each TSA-seq high-slope center; 2) The superposition of different groups of TSA-seq high-slope centers with varying positions of active chromatin features, including two groups corresponding to active marks more enriched on either the left or the right side of the high-slope center.

Cluster analysis of high-slope domains confirms the second explanation (Fig. 5B and Supplemental Fig. S9). Using the H3K4me1 mark for clustering, we observed distinct patterns of H3K4me1 enrichment in each of the 6 clusters, with patterns of H3K4me3, H3k27ac, and H3K9ac in each cluster similar to the H3K4me1 pattern in that same cluster. All 6 clusters contained peaks of active transcription marks centered at ∼100-125 kbp to one side or the other of the TSA-seq high-slope centers. These peaks of transcription-related marks each appeared ∼100 kbp in width (full-width, half-peak). The two largest clusters (C1, C3) show pronounced peaks of active marks plus an overall lower than peak-level enrichment of the active mark on the same side of the peak with an accompanying depletion of the active mark, and enrichment of the repressive H3K9me3 mark, on the opposite flanking region (Fig. 5B). Two clusters (C2,C6) showed less extension of these lower levels of active marks distal to the peak at ∼125 kb from the DLCD center. The remaining two clusters (C4, C5) showed low levels of enrichment of active marks, relative to peak levels, but on the contralateral side of the DLCD center.

Overall, DLCDs mostly map to the boundary of active versus repressive genomic domains and contain a peak of active mark enrichment at ∼125 kb to one side of the DLCD (Fig. 5A-B).

We applied the same enrichment and cluster analysis to the analysis of the distributions of histone modifications, transcription-related factors (“TFs”), and chromatin structural proteins relative to Hi-C TAD boundaries. Most of these marks showed sharper, but occasionally broad, unimodal enrichment distributions in the averaged heat-maps (Supplemental Fig. S10). Cluster analysis revealed almost half of the TAD boundary clusters showed a similar asymmetric distribution of marks to one side of the TAD boundaries as we observed for distributions of the same marks relative to TSA-seq high-slope centers, although without the consistent prominent peaks at ∼100 to one or the other side of the boundary (Supplemental Fig. S10). However, some major clusters instead show symmetric distributions of these marks relative to the TAD boundaries, resulting in the overall unimodal distributions observed in the profile plots (Supplemental Fig. S10). This is more evident in H3K4me1 and H3K27ac profile plots, in which remnants of the bimodal distribution appear flanking the central peak (Supplemental Fig. S10). Thus, a higher fraction of TSA-seq high-slope centers versus TAD boundaries define the boundaries of active and repressed chromatin domains.

The cluster analysis of TAD boundaries does reveal a narrow center peak enrichment of CTCF and SMC3 at most TAD boundaries, consistent with previous profile plots (Dixon et al. 2012), but this is in addition to their broad enrichment to one side of the TAD boundary for a number of these clusters (Supplemental Fig. S11).

### DLCDs are enriched in transcription factors, transcriptional coactivators, and subunits of chromatin remodeling and modifying complexes, including those related to acidic activators

We calculated enrichment ratios of the various factors mapped by ChIP-seq, dividing the height of the peak by the genomic background in the enrichment heatmaps, and sorted them based on this enrichment ratio (Fig. 5C, Supplemental Fig. S8). A large number of transcription factors, transcriptional activators, and chromatin remodeling and modifying complex subunits show enrichment over DLCDs in very similar patterns to those described in the previous section for TSSs, super-enhancers, POL2RA, H3K4me3, H3K4me1, H3K9ac, and H3K27ac.

Notably, the 5 factors with the highest enrichment ratios all show connections with acidic activation transcription factors (e.g. VP16 and NF-kB). RELA (p65) was 3^rd^ on this list of factors and is the transcriptional activator subunit of NF-kB, (Schmitz and Baeuerle 1991). The top sorted factor, NSD2, directly binds to NF-kB (Yang et al. 2012) and has H3K36 methyltransferase activity, a histone modification associated with active genes bodies. ZZZ3, the number 2 sorted factor, is a subunit of the Ada-two-A-containing (ATAC) histone acetyltransferase complex and contains the ZZ H3-reader domain (Mi et al. 2018). ZZZ3 binding sites are enriched in DNA binding motifs for CREB1, an acidic transcriptional activator (Krebs et al. 2011). SAGA, another, GCN5-containing HAT containing a different Ada2 protein, is recruited early by the VP16 acidic activator domain (Memedula and Belmont 2003) and considered an essential target for gene activation by the Gal4 acidic activator (Bhaumik and Green 2001). Although still poorly characterized relative to SAGA, the ATAC complex is thought to recruit similar coactivators as SAGA and may have similar important functions for acidic activator function.

CREB3, ranked 4^th^ on the list, is another transcriptional activator containing an acidic activator domain. CREB3 binds to HCF1 (Lu et al. 1997), the endogenous protein-binding partner of VP16. HCF1 is also a binding factor for myc (Thomas et al. 2016) and a promoter of cell-cycle progression (Mangone et al. 2010).

BRD4, 5^th^ on the list, is a histone reader and transcriptional regulator recruited to enhancers and especially super-enhancers (Loven et al. 2013). BRD4 binds p-TEFb (Itzen et al. 2014) as well as binding and/or being recruited by multiple transcription factors, including the acidic transcriptional activators p53, RELA (Huang et al. 2009; Wu et al. 2013), and VP16 (Zhao et al. 2011).

## Discussion

### Identification of DLCDs

Here we demonstrated the use of TSA-seq, which reports on microscopic distance to particular nuclear structures, to identify chromosome regions with relatively high levels of large-scale chromatin decondensation. How rapidly the TSA-seq score changes as a function of genomic distance will be a function of both the local chromosome compaction and the chromosome orientation relative to the staining target. More specifically, TSA-seq score high-slope regions should detect decondensed chromosome regions in chromosome trajectories oriented more perpendicular, rather than parallel, to the TSA staining targets. The slope of the TSA-seq score depends on the orientation of a chromosome region relative to the staining target as well as its large-scale chromatin compaction. Despite this, many high-slope regions (top decile of slope magnitude) identified using either anti-speckle or anti-lamin TSA staining are the same, although some are specific to one or the other TSA staining target. Indeed, we primarily detect DLCDs using SON and lamin TSA-seq in the transition zones connecting TSA-seq peaks with valleys. Previously we demonstrated that in K562 cells these transition zones corresponded to approximately linear interphase chromosome trajectories running between the nuclear lamina and nuclear speckles, as visualized by 3D immuno-FISH. We do not expect our approach to be able to detect DLCDs, or other types of decondensed chromatin domains, lying in chromosome trajectories running parallel to the TSA staining target. For example, SON TSA-seq peaks correspond to genomic regions with a very high density of highly expressing genes, super-enhancers, and histone marks associated with active chromatin (Chen et al. 2018; Wang et al. 2021); these peaks are likely to include decondensed chromatin domains, but they are all similarly close to speckles and thus appear as regions of low TSA-seq slope.

Therefore, we assume DLCDs detected as TSA-seq high-slope regions represent a subset of the total number of decondensed large-scale chromatin regions. Likewise, we also do not expect all DLCDs detected using one TSA staining target, such as nuclear speckles, to appear in TSA-seq directed against a different staining target, such as lamins. Significantly, though, we did find a large degree of overlap between DLCDs detected as high-slope regions in TSA-seq data sets against different targets.

We see our method as primarily useful to recognize a class of unusually decondensed large-scale chromatin domains, given that the derivative of the TSA-seq is noisy and highly reproducible only for the top 10% of slope magnitudes.

Importantly, we observed a high degree of overlap between DLCDs detected as TSA-seq high-slope regions and DLCDs detected as Hi-C compartment score (EV1) high-slope regions. While the TSA-seq derivative can be mathematically related to changes in distance traversed by a chromosome region, and therefore chromatin compaction, the Hi-C PC1 slope is more difficult to interpret. To the degree that Hi-C compartments are related to the colocalization of chromosome regions in the same nuclear regions, heuristically, the EV1 slope should reflect how fast a chromosome region traverses from one nuclear location to another and therefore chromosome compaction. Here, however, by demonstrating the empirical correlation between TSA-seq and Hi-C PC1 high-slope domains, we better justify the detection of DLCDs from Hi-C PC1 data as a valid alternative when multiple TSA-seq datasets are not available.

### DLCD Features and Connections with Acidic Transcriptional Activators

The first salient feature of DLCDs is their detection near the subset of Hi-C topological domain boundaries corresponding to a large extent compartment or subcompartment boundaries and the boundaries separating active from inactive chromatin domains. Their second salient feature is that they contain ∼100 kb-wide peaks of multiple genomic features and chromatin marks associated with active transcription, including chromatin modifying and remodeling complexes, which appears distinctive for this subset of Hi-C topological domain boundaries near DLCDs, as compared to peaks of CTCF and cohesin that are seen for all TAD boundaries (ref, Supplemental Fig. S12).

What remains unclear is why these peaks of active marks near the DLCDs are displaced by ∼100 kb from the CTCF/cohesin peak closer to the DLCD center. As mentioned in Results, use of a larger smoothing window for our calculation of the 3^rd^ order polynomial fitting the TSA-seq data reduces noise but may increase a systematic bias in shifting the DLCD center position. Given the known linkage between transcription-related factors and chromatin decodensation, we suspect that the true center of decondensation of the DLCDs therefore might be these active mark peaks, which are also the main drivers for the observed large-scale chromatin decondensation.

Indeed, this is why we are so intrigued by super-enhancers having a pronounced enrichment in DLCDs, as well as multiple transcription-related factors in our top-sorted peak enrichment list which are related to acidic transcriptional activators. Twenty years ago, our laboratory demonstrated an ∼30-fold extension of a large, heterochromatic gene-amplified chromosome region after tethering the VP16 acidic activation domain to this region via a lac repressor fusion protein (Tumbar et al. 1999). This large-scale chromatin decondensation activity after tethering was a common property of all tested acidic activator domains, including from VP16, Gal4, p65 (RELA), and p53, but not several other classes of transcriptional activators (Carpenter et al. 2005). Tethering acidic activation domains rapidly recruited components of SAGA, SWI/SNF BRG1 and BRM complexes,p300/CBP, PCAF, and TIP60 (Memedula and Belmont 2003). Using a similar tethering assay, we also showed that both VP16 and p65 (RELA) acidic activation domains induced long-range chromosome movement from the nuclear periphery to interior (Chuang et al. 2006).

Super-enhancers are defined by high concentrations of Mediator and p300/CBP, both recruited by acidic activators. Of the 5 factors with the highest ChIP-seq enrichment near DLCDs, CREB3 and RELA (p65), a NF-kB subunit, are both acidic transcriptional activators, and CREB3 is known to bind HCF1, a cellular factor that is also recruited by the viral VP16 acidic activator, As discussed in more detail in Results, the remaining three proteins on the list-NSD2, ZZZ3, BRD4-are either known to be recruited by acidic activators, or are in chromatin modifying complexes closely related to other chromatin modifying complexes known to be recruited by acidic activators. Moreover, BRD4, recruited by multiple transcription factors including acidic activators, is also enriched at super-enhancers. Indeed, BRD4 tethered to plasmid transgene arrays induces large-scale chromatin decondensation similar to that demonstrated by tethering acidic activators (Zhao et al. 2011). NSD2, #1 on our top-sorted list, changes TAD boundary strengths (61 increased, 5 decreased) and A/B compartmentalization when overexpressed in multiple myeloma as a result of the t(4;14) chromosome translocation (Lhoumaud et al. 2019).

### Working Model and Future Directions

In conclusion, our results suggest a working model in which DLCDs represent chromatin regions of high large-scale decondensation that are located adjacent to boundaries separating transcriptionally active from inactive chromatin domains, which frequently correspond to Hi-C compartment or subcompartment boundaries (Fig. 6B, blue). There may be additional chromatin regions with comparable or even higher levels of large-scale chromatin decondensation that are not detected by the current SON and lamin TSA-seq datasets due to their roughly parallel orientation relative to nuclear speckles or nuclear lamina, respectively. Indeed, as illustrated in Fig. 6B (orange DCLDs), we imagine the possibility of similar decondensed regions near nuclear speckles that would appear as SON TSA-seq low-slope regions, as well as near other possible nuclear compartments (Fig. 6B, arrowhead). Likewise, given the limited reproducibility of TSA-seq slope measurements at lower slope magnitudes, we cannot exclude the existence of decondensed large-scale chromatin regions at other TAD boundaries; if present, however, our results would suggest these chromatin regions are not quite as decondensed and/or as large as DLCDs.

**Figure 6.**
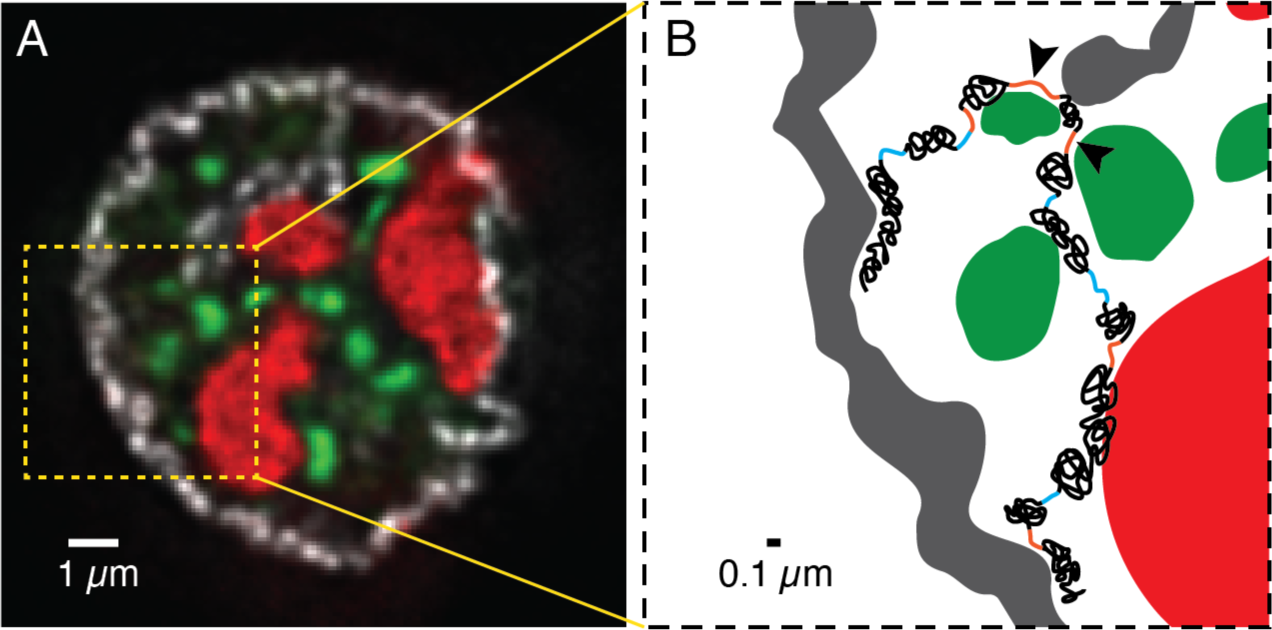
DLCDs detected by high-slope TSA-seq regions separate active and inactive chromatin domains. **(A)** Immunofluorescence deconvolved image (single optical section) of a typical K562 nuclei. Nuclear lamina (grey), nucleoli (red), nuclear speckles (green) are stained with mouse-anti-RL1, anti-MKI67IP-CF488A, and anti-SON-CF647 respectively. Scale-bar = 1 µm. The yellow dashed box is enlarged for **(B),** a schematic model of a local chromosome trajectory. Decondensed large-scale chromatin domains (DLCDs) oriented in trajectories roughly perpendicular to the TSA-seq staining target will appear as high-slope regions in the TSA-seq score. The binary images of nuclear lamina (grey), nucleoli (red), nuclear speckles (green) are produced from A, with added regions of compact chromatin (black), high-slope DLCDs (blue), and decondensed large-scale chromatin domains located in low-slope TSA-seq regions (orange). Chromosome trajectories, even if decondensed, that run parallel to the TSA staining target compartment will not be detected as high-slope in the TSA-seq score (arrowhead). Scale-bar is 0.1 µm.

Based on Hi-C experiments in the absence of cohesion complex, loop extrusion and compartmentalization have been proposed as major chromatin organization mechanisms (Schwarzer et al. 2017; Rao et al. 2017). DLCDs separated domains with active vs. repressive chromatin marks and located near a subset of Hi-C domain boundaries (compartment, subcompartment, and TAD). Thus, DLCDs may enhance the genome compartmentalization by spatial separation of active and inactive portions of the genome.

Moving forward, several questions await further study. First, what is the exact ultrastructure of DLCDs? We are intrigued by the possibility that they might correspond to the very decondensed chromatin regions connecting large-scale chromatin domains, as previously visualized by electron microscopy (see figure 6 from (Belmont and Bruce 1994)). Second, what are the dynamics of these DLCDs? We showed that only ∼50% of the regions we visualized by 3D FISH showed noticeable decondensation by wide-field microscopy, suggesting DLCD chromatin compaction is dynamic. The timescale for such dynamics would have large implications for the possible physiological significance of DLCDs. Finally, what are the functional consequences of deleting DLCDs? Previous work has revealed relatively subtle effects on gene expression after knockdown of cohesin at Hi-C domain boundaries (Schwarzer et al. 2017; Rao et al. 2017). DLCDs are distinct among the larger class of Hi-C domain boundaries in having flanking peaks of transcription-related marks and genomic features. DLCD deletion experiments will be critical in revealing possible DLCD roles on maintaining normal levels or stability of gene expression in their flanking chromatin domains.

## Methods

### Genomic Datasets

TSA-seq score and normalized Hi-C interaction matrix were downloaded from the 4D Nucleome consortium data portal (https://data.4dnucleome.org/). Histone modification ChIP-Seq enrichment over input datasets were downloaded from the ENCODE consortium (https://www.encodeproject.org/). Transcription factor ChIP-Seq peaks datasets were downloaded from the ReMap catalogue (http://tagc.univ-mrs.fr/remap/index.php) (Chèneby et al. 2020). See Supplemental Table S3 for the complete list of datasets used.

### Measuring TSA-seq derivative

TSA-seq score datasets were previously calculated in 20 kb bins. For each bin, a cubic polynomial was fit to the data corresponding to a window of 2N +1 bins centered at this bin using “lm” function from R stats package (R Core Team 2020) with the raw option on. The polynomial was used to calculate a smoothed TSA-seq score, and the first-order coefficient of the polynomial was saved as the estimate for the TSA-seq score slope. If the TSA-seq score value was missing from any bin in a given window, the slope values were recorded as zero for the central bin of this window. This included using zero values for regions less than N bins from the chromosome ends or from unmapped chromosome regions. After comparison of results with different size windows, a window size of 21 bins was chosen for all subsequent analysis.

### Identifying high-slope domains and domain centers

High-slope domains called from individual TSA-seq datasets were identified as contiguous bins, each with absolute value first derivative values in the highest decile (top 10 percentile). The center of a high-slope domain (high-slope center) was identified as the bin with the highest absolute magnitude of the slope (i.e. conceptually where the second derivative should approach zero).

To create a reference set of high-slope domains called from multiple TSA-seq datasets, two strategies were used. 1) A mean TSA-seq slope was calculated by averaging the absolute value of all the available TSA-seq slopes (non-normalized raw values) in K562 from each dataset in every genomic bin. High-slope domains were then identified as described above for individual TSA-seq datasets but using the highest decile of this mean slope dataset. 2) Majority high-slope domains were identified by a voter scheme strategy. A genomic bin was considered high-slope if a simple majority of TSA-seq datasets identified it as high-slope (top decile). For both mean and majority high-slope reference tracks, we first merged the domains that were separated by 20 kb (1bin) and counted only the domains 40kb (2 bins) or larger in length.

To compare the percent overlap between different TSA-seq high-slope datasets we used the Jaccard index (Fig. 1F and Supplemental Fig. S3). The overlap between dataset A and B was defined as the average of *length(A∩B)/length(A)* and *length(B∩A)/length(B)*.

### Cell Culture

K562 cells were obtained from the ATCC and cultured according to the ENCODE Consortium recommended protocol (http://genome.ucsc.edu/ENCODE/protocols/cell/human/K562_protocol.pdf). Briefly, K562 cells were cultured in RPMI-1640 medium supplemented with 10% FBS (Sigma cat#: F8067) and 1%V/V 100x Antibiotic-Antimycotic (Gibco) and passaged/harvested at 0.8 106 cells/mL. HCT116 cells were also obtained through the ATCC and cultured according to the 4DN Consortium recommended protocol (https://data.4dnucleome.org/resources/experimental-resources/cell-lines). Briefly, HCT116 cells were cultured in McCoy’s 5A medium supplemented with 10% FBS (Sigma cat#: F8067).

### DNA 3D Fluorescence In Situ Hybridization (FISH)

FISH probes were prepared using BACs (Invitrogen). BACs were extracted from bacterial culture using the NucleoBond® Xtra Midi plasmid extraction kit (Macherey-Nagel) according to the manufacturer’s provided instructions. 5 µg of each BAC was fragmented with a restriction enzyme cocktail (5U of AluI, DpnI, HaeIII, MseI, MspI, and RsaI from NEB) to achieve fragment sizes of ∼150bp. This fragmented BAC DNA was labeled as described elsewhere (Dernburg 2011) using either Biotin-14-dATP (Invitrogen) or Digoxigenin (Dig)-11-UTP (Roche) using Terminal Deoxynucleotidyl Transferase (TdT; Thermo Scientific) with the following reagent composition: fragmented BAC 2 µg, TdT 12 U, TdT buffer 1X, unlabelled nucleotide (dATP or dTTP; NEB) 108 µM, and labeled nucleotide (Dig or Biotin) 54 µM.

Our 3D-FISH procedure was modified from previously described protocols (Markaki et al. 2013; Solovei and Cremer 2010). For K562 cells, our 3D-FISH procedure is as described previously (Chen et al. 2018) with the following minor changes. To enhance K562 cell attachment to coverslips, prior to cell attachment coverslips were treated with 0.1 % W/V poly-L-lysine (mol wt 70,000–150,000 molecular weight; Sigma-Aldrich) for 60 mins at room temperature (RT), excess liquid was removed, and coverslips dried overnight. Alternatively, coverslips were treated with 0.1 % W/V poly-L-lysine (mol wt 150,000-300,000 kDa; Sigma-Aldrich) for 60 mins at RT, rinsed twice with distilled water, and dried overnight. The K562 cell suspension in complete growth media was added to round 12 mm #1½ glass coverslips (0.5mL at 5×10^5^cells/mL) and incubated for 45 minutes in a humidified incubator with 5% CO2 at 37°C.

3D-FISH with EdU labeling was done in HCT116 cells (data related to Supplemental Fig. S5). HCT116 cultures at ∼70% confluency were trypsinized and 5×10^5^ cells (in 0.5mL of growth media) were seeded per well (24-well plate; the volume for all the wash/rinse steps was 500µl); each well contained a round 12 mm #1 ½ glass coverslip. After two days of cell culture, cells were pulse-labeled with growth media containing 5µM EdU (Invitrogen) for 15 mins. Cells were rinsed once with 37°C calcium, magnesium free phosphate buffered saline(CMF-PBS; KCl 2.67 mM, KH2PO4 1.47 mM, NaCl 136.9 mM, Na2HPO4 8.1 mM) and fixed with freshly prepared 3% PFA (paraformaldehyde) in CMF-PBS (pH 7.4) for 10 mins at RT and then free aldehydes were quenched with 0.5 M glycine in 1X CMF-PBS for 5 mins at RT. Coverslips were rinsed 3 times (3x) with CMF-PBS and cells permeabilized with 0.5% Triton X-100 (Thermo Scientific, catalog #28314) in CMF-PBS for 10 mins at RT. Cells were then rinsed 3x with CMF-PBS and incubated in 20% glycerol/CMF-PBS at RT for 1 hour. Alternatively, cells were incubated in 20% glycerol/CMF-PBS at 4°C overnight. Coverslips were frozen in liquid nitrogen and thawed on a paper towel at RT for a total of 4 repetitions, washed 3 × 10 mins with CMF-PBS at RT, rinsed in 0.1 N HCl, and then incubated in 0.1N HCl for 10 mins at RT. Coverslips were then rinsed in 2X-SSC (1X SSC: 150mM sodium chloride, 15mM sodium citrate) 3x and incubated for >24 hrs in 50% formamide (Sigma, catalog # 47671) / 2X-SSC (pH 7.0) at 4°C prior to hybridization. We observed a variable increase in FISH signal area that appeared to correlate with increased storage time in formamide; therefore we always included an internal control FISH for comparison with the test FISH probe signals.

40-80 ng of each probe (labeled with biotin or Dig) and 1µg of unlabeled human COT1-DNA (Invitrogen) were dissolved in 4µl of hybridization buffer (10% dextran sulfate, 50% formamide, and 2X-SSC) and mixed by vortexing. Per 12 mm round coverslip, 4 µl of the probe mixture was added and then coverslips were sealed onto glass slides with rubber cement (Elmer’s). Slides were placed on a hot plate at 76°C for 3 mins to denature the DNA and then hybridized for 72 hrs in a humid chamber at 37°C.

After hybridization, coverslips were floated by adding several drops of 2XSSC and gently removed to avoid shearing of cells. Coverslips were washed at 37°C 3 × 5 mins in 2X-SSC, then at 60°C for 3 × 5 mins in 0.1X-SSC, and finally at RT with 4X-SSC for 5 mins. Coverslips were washed with blocking buffer (4X-SSC/0.1% Triton X-100/4% BSA (Bovine serum albumin; Sigma, catalog #A7906)) at RT for 30 mins. Coverslips were then incubated in the dark for 2 hrs with streptavidin–Alexa Fluor 594 (Jackson ImmunoResearch Laboratories) and mouse anti-Dig–Alexa Fluor 647 (Jackson ImmunoResearch Laboratories) diluted 1:200 in blocking buffer and then washed in 4X-SSC/0.1%Triton X-100 3 × 5 mins.

EdU then was reacted with Alexa-488-azide (Click Chemistry Tools) using the Click-iT™ EdU Cell Proliferation Kit for Imaging (Invitrogen) according to the manufacturer’s provided instructions.

Coverslips after either K562 or HCT116 FISH were mounted on glass slides (Fisher Scientific) with DAPI-containing antifade mounting media (0.3 μg/ml DAPI (SigmaAldrich)/10% w/v Mowiol 4-88(EMD Millipore)/1% w/v DABCO (Sigma-Aldrich)/25% glycerol/0.1 M Tris, pH 8.5).

### Microscopy and data analysis of FISH experiments

Microscopy Image acquisition was done using the OMX-V4 microscope (GE Healthcare) equipped with a U Plan S-Apo 100×/1.40-NA oil-immersion objective (Olympus), two Evolve EMCCD cameras (Photometrics). Z-sections were 200 nm apart. For each 3D-FISH condition n≥100 cells per coverslip (two technical replicate coverslips) were imaged per experimental condition.

We used intensity thresholding to measure FISH signal projected areas. 16-bit imaging files were imported into python using a python wrapper for Bio-Formats (McQuin et al. 2018;

Linkert et al. 2010). Each FISH spot was cropped using a 24×24×11(X×Y×Z) voxel volume centered around the brightest pixel of the FISH signal. Background noise was estimated using the mode of the intensity histogram computed in this cropped volume; because the FISH signal volume was small relative to the total cropped box volume, this histogram mode approximated the mean background intensity. The background mode intensity was subtracted at each pixel and then the volume was projected onto the x-y plane using the sum intensity at each pixel along the z-dimension. This projection was segmented using a threshold corresponding to 50% of the peak intensity to produce a binary image.

We used the Feret diameter as a length measure of the FISH signal. The Feret (caliper) diameter is defined as the longest straight line that connects two points on the edge of a convex shape. For the 3D-FISH experiment in which we controlled for cell cycle phase, we manually assigned each FISH signal as corresponding to either a G1/G2 (no EdU foci) or S-phase (EdU replication foci visible) nucleus. A two-sided Welch two-sample t-test was used to test the hypothesis that shared and test probes have equal means.

### EV decomposition of HiC intrachromosomal contacts

To identify K562 A/B compartments and compute the related Hi-C EV1, HiC intrachromosomal contacts were calculated at different resolutions using Cooltools (Venev et al. 2020). To assign the correct sign (+/-) for EV1 values, the genome-wide GC content was used as a proxy to match the A compartment with EV1 positive values. A and B compartments were assigned as the positive and negative values of the EV1, respectively, using the EV1 calculated at 100kb resolution. For EV1 slope calculations, the EV1 was calculated at 10 kb resolution and binned at 20 kb to match TSA-seq slope bins. Finally, the EV1 slope was calculated exactly as described for the TSA-seq slope.

### Identifying chromatin organization domain boundaries

K562 Hi-C TAD boundaries (arrowhead TAD annotations (Rao et al. 2014)) were downloaded from GEO and lifted from hg19 to hg38 using the UCSC LiftOver tool (https://genome.ucsc.edu/cgi-bin/hgLiftOver).

K562 Hi-C subcompartment calls in hg38 were imputed using SNIPER, a machine-learning approach that uses Hi-C subcompartment calls from cell lines with high-read Hi-C data to impute Hi-C subcompartments in cell types with sparser Hi-C data (Xiong and Ma 2019).

K562 A/B compartment boundaries were identified as the boundary of the two consecutive bins were the EV1 signal crosses the y=0 line, using the EV1 calculated at 100 kb resolution. We defined the uncertainty of all domain boundary calls as +/- 1/2 the resolution of the relevant dataset. If the boundaries of two domains were closer than this resolution to each other, these two domains were merged into one domain.

The observed over expected (obs/exp) overlap probabilities for Hi-C domain boundaries, LAD/iLAD boundaries, TSA-seq high-slope regions, and EV1 high-slope regions were calculated using *GenometriCorrelation* function from GenometriCorr package (Favorov et al. 2012).

### Transcription factor, histone modification, and genic feature profile plots

We used ChIP-Seq and DNase 1-seq datasets from ENCODE. For ChIP-seq of histone modifications and RNApol2, we used pulldown versus input ratios for our data analysis. For ChIP-seq of transcription factors, transcriptional coactivators, and subunits of chromatin remodeling and modifying complexes, we used called peaks from ReMap (Chèneby et al. 2017). For DNase 1-seq datasets, we used read-depth normalized signal from ENCODE. Both pulldown/input and read-depth normalized signal were obtained using the ENCODE standard pipeline mapped to hg38. For transcription factor ChIP-Seq we used called peaks from ReMap (Chèneby et al. 2017). For K562-specific super-enhancer and enhancer annotations, we used data from SEdb (Jiang et al. 2019). List of genes for hg38 were obtained from Bioconductor (Bioconductor Core Team and Bioconductor Package Maintainer 2019) and transcription start sites were identified as the 5’ of the genes.

For pulldown/input measurements, average values were computed for every 10 kb bin in the genome. For peak count measurements, enhancers, and transcription start sites, the feature count was calculated for every 10 kb in the genome. In cases that more than one dataset was available for called peaks of a particular TF, datasets were consolidated using the following majority voter scheme. A called peak was considered a true call if it was called in the simple majority of available datasets. For generating profile plots and heatmaps, the signal was calculated ±500kb in 10kb bins (total of 101 bins) for every high-slope center. The data was either summed for profile plots or were stored as an m×101 array (where m is the number of high-slope domains) for heatmaps.

### Plotting heatmaps

For plotting the heatmaps, we used hierarchical clustering to cluster high-slope domains feature matrix. From the m×101 heatmap array, a distance matrix was created by computing the pairwise 1-*ρ*(m_i_,m_j_), where *ρ*(m_i_,m_j_) is the Spearman correlation of high-slope domain i and j of the feature matrix. This distance matrix was then used as a metric in the hierarchical clustering. Hierarchical clustering was done using hclust function of R stats package using the Ward’s minimum variance method (ward.D2 option). For some heatmaps, as noted, in each row the feature matrix was normalized by computing the z-scores of each 10 kb bin. Heatmaps were visualized using pheatmap package (Kolde 2019).

### Software Availability

Source code used for calculating slopes, analyzing data, and generating all figures in this paper is available at https://github.com/belmontlab/gholamalamdari_etal_2021. R Markdown notebooks available at this site contain additional details of methods used in all computational analysis.

## Acknowledgments

We thank Drs. William Brieher, Lisa Stubbs, and Brian Freeman (UIUC) for helpful suggestions during the course of this research. We thank members of the Belmont U54 NIH Center-Drs. Jian Ma, David Gilbert, Bas van Steensel and members of their laboratories-for discussions and helpful suggestions. This work was supported by National Institutes of Health grant R01 GM58460 to A.S. Belmont. The authors declare no competing financial interests.

## Author contributions

Computational tool development to estimate TSA-seq slope, and quantitative analysis of TSA-seq slope data and integration with other genomic data were done by OG with guidance from ASB. FISH validation experiments and the development of quantitative image analysis pipeline were done by OG with guidance from ASB. YC, OG, LZ, and ASB conceived the study and LZ performed exploratory FISH experiments in NIH-3T3 cells. The manuscript was written by OG and ASB.

