## Supplementary Figures for "Identification of decondensed large-scale chromatin regions by TSA-seq and their localization to a subset of chromatin domain boundaries"

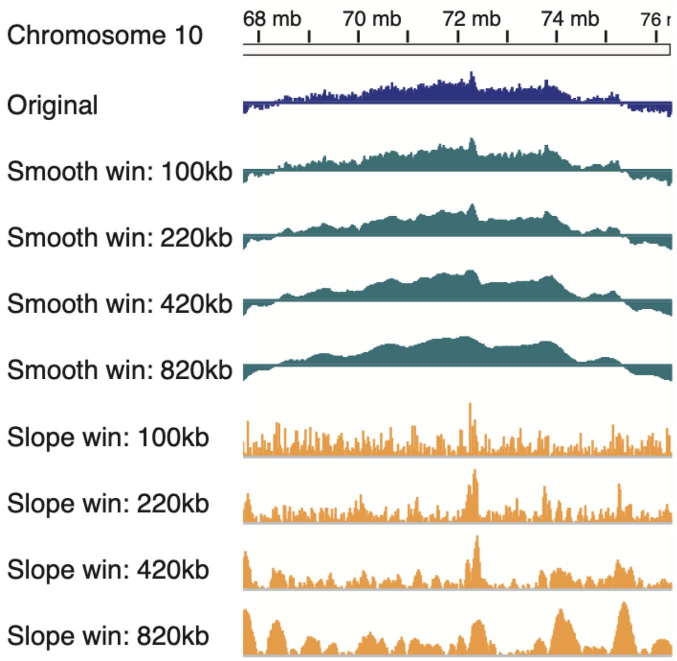


**Supplemental Figure S1 | TSA-seq score smoothing and estimation of slope using different window sizes for fitting the TSA-seq score to a cubic polynomial.** The original data is mapped using 20 kb bins. This data was then approximated by fitting a cubic polynomial within a window centered around each bin. Windows of 5, 11, 21, and 41 bins were used corresponding to sizes of 100, 220, 420, and 820 kb. Tracks shown for a region of Chr10 from top to bottom: SON TSA-seq score replicate 1 technical replicate 1, TSA-seq score smoothed by fitting to windows of 100, 220, 420, and 820 kb, and absolute values of derivatives calculated from polynomial fit using windows of 100, 220, 420, and 820 kb. For further analysis, we chose a window size of 420 kb as a compromise between minimizing noise and maximizing spatial resolution.

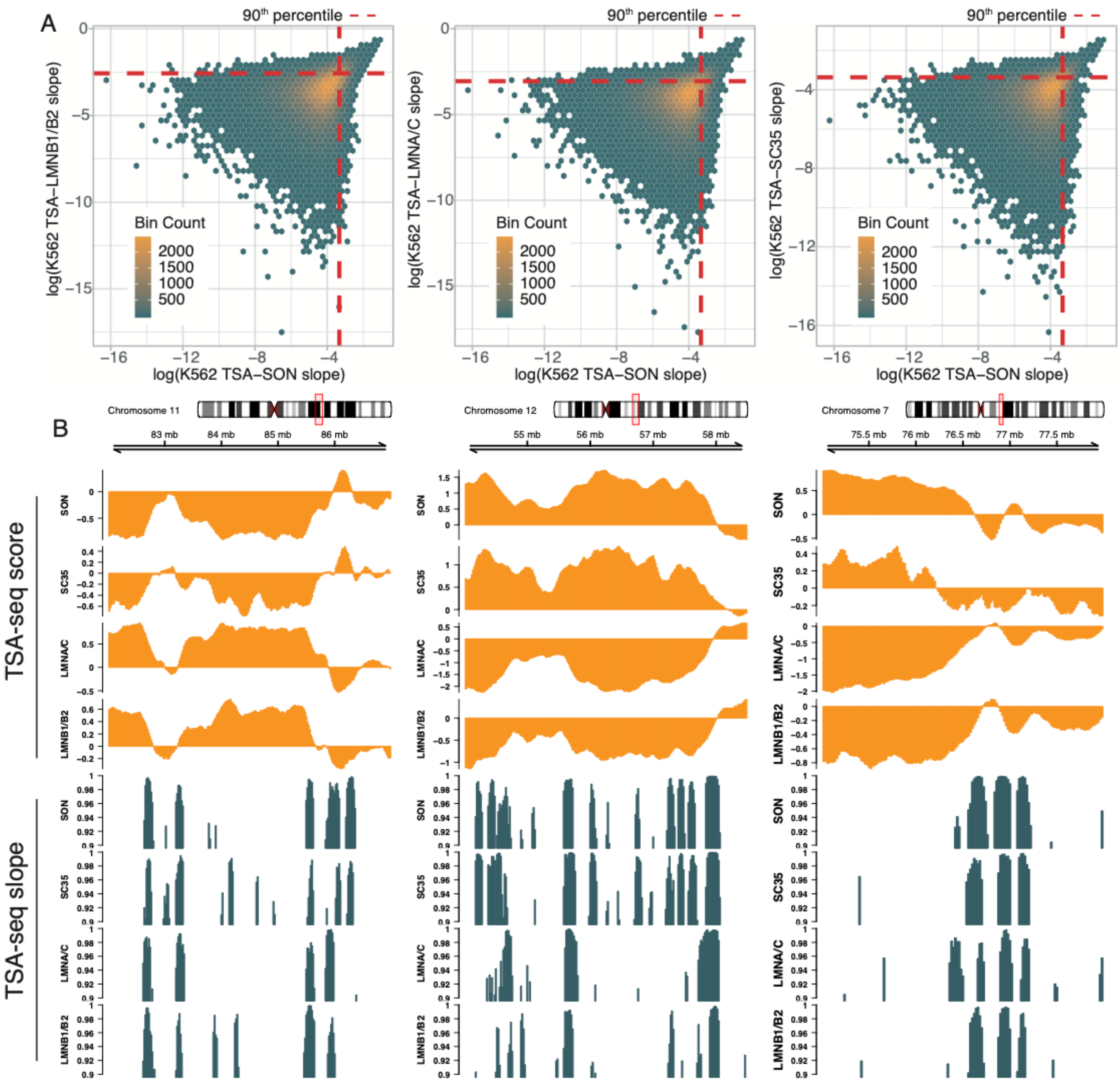


**Supplemental Figure S2 | TSA-seq high-slope regions calculated from TSA-seq datasets produced using different staining targets show extensive overlap.** The slope of the TSA-seq score depends on the orientation of a chromosome region relative to the staining target as well as its large-scale chromatin compaction. Despite this, many high-slope regions (top decile of slope magnitude) identified using either anti-speckle or anti-lamin TSA staining are the same, although some are specific to one or the other TSA staining target. **(A)** Binned scatter plots of TSA-seq slopes calculated from two different TSA-seq datasets showing the number of bins with given values of the natural log (from left to right) using SON versus LMNB1/2, SON versus LMNA/C, or SON versus SC35 TSA-seq. Slopes correlate well for higher slope values and especially for the top decile of slope values (red dotted lines mark 90%). Nearly comparable correlations in slopes are observed comparing lamin versus speckle TSA-seq datasets as observed comparing TSA-seq datasets corresponding to different speckle markers (SON, SC35). **(B)** Comparison of high-slope domains (top decile) identified from TSA-seq against nuclear speckles (SON or SC35) and nuclear lamina (LaminB1/B2 or LaminA/C). Top to bottom: chromosme ideograms, smoothed SON, SC35, LMNB1/B2, and LMNA/C TSA-Seq scores, and slopes (top decile) calculated from SON, SC35, LMNB1/B2, and LMNA/C TSA-seq datasets.

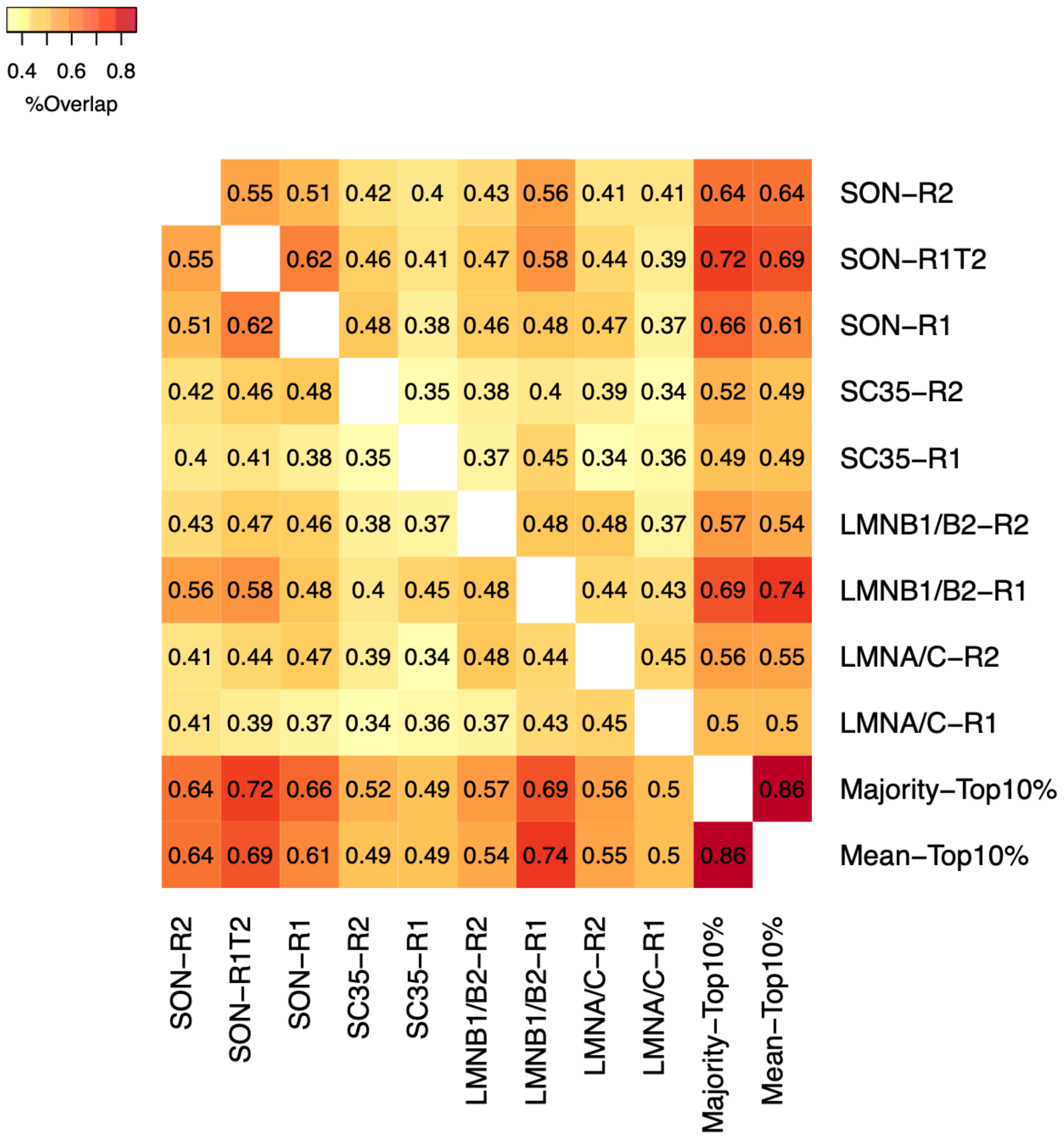


**Supplemental Figure S3 |** Percent overlap (Jaccard index) between high-slope domains identified using individual TSA-seq datasets with merged high-slope domains (mean and majority). For each target there are two biological replicates (R1 and R2). For SON-R1, T2 denotes technical replicate 2 of the SON TSA-seq. High-slope domains were consolidated using two approach: 1) Mean- Averaging slopes from each TSA-seq dataset to calculate a mean slope at each bin, and then identifying the top 10% values of this mean across the genome; 2) Majority- including all high-slope domains that were identified as high-slope domains in the majority of TSA-seq individual datasets. The Jaccard index is defined as the ratio of the intersection to union of two sets of elements. See Methods for implementing the calculation of this index for high-slope domains.

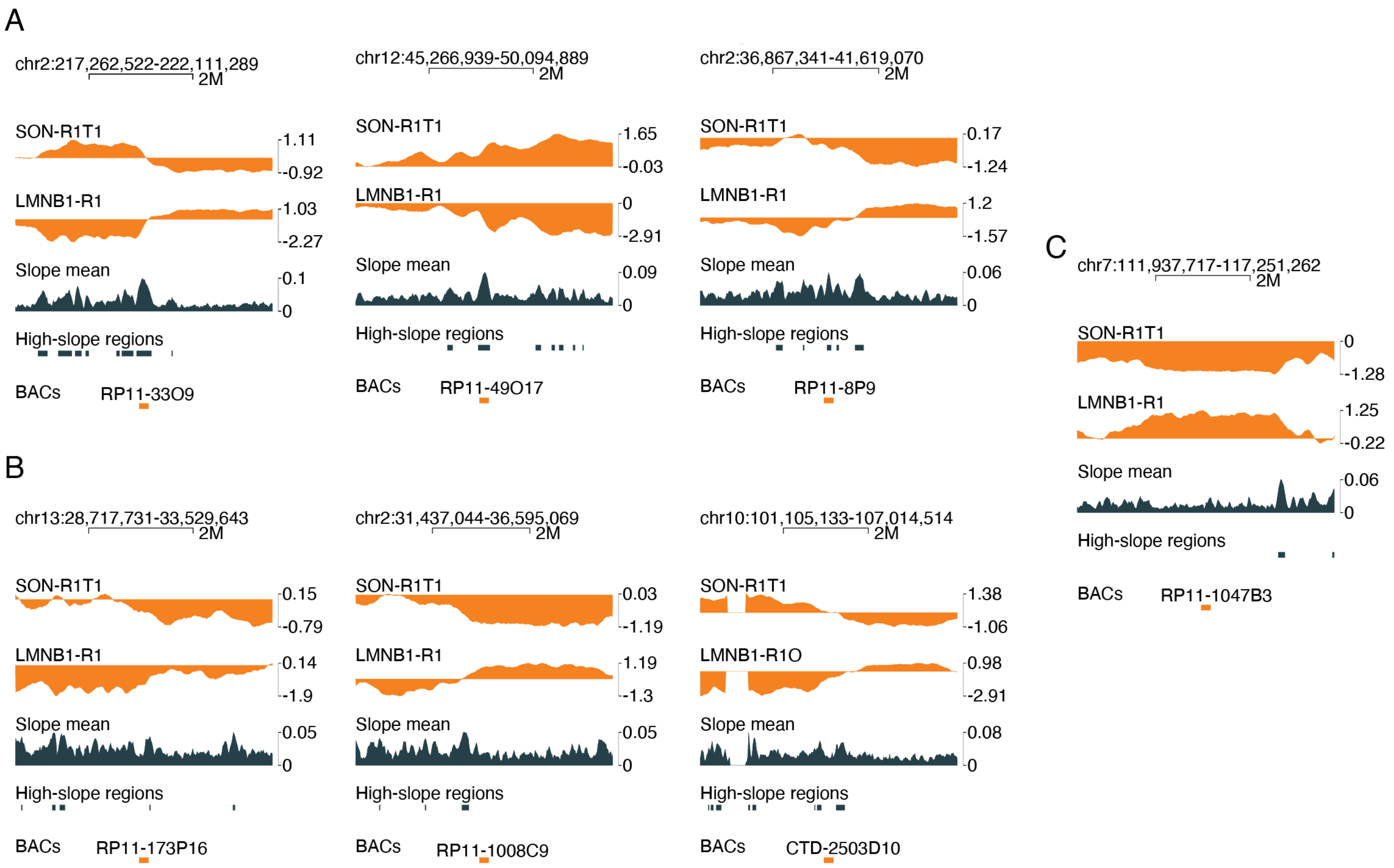


**Supplemental Figure S4 Genomic location of BAC probes used in this study.** (A) Probes from high-slope regions (A), probes from low-slope regions (B), and the shared-probe used as the internal control for FISH experiments (C). Top to bottom for each genomic region: genomic coordinates, smoothed SON and LMNB1/B2 TSA-Seq scores, average of TSA-seq slopes calculated across all available TSA-seq slope datasets (Slope mean), and high-slope domains identified based on top decile of the mean of all TSA-seq score slopes (High-slope regions).

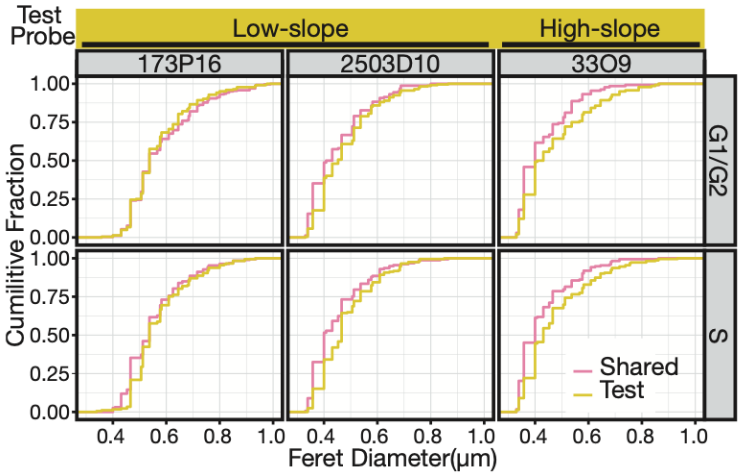


**Supplemental Figure S5 | DLCD decondensation is independent of cell cycle in HCT116 cells.** Cumulative distribution of the 3D-FISH signal Feret diameter for cells in G1/G2 versus S. The G1/G2 population is dominated by cells in G1 (~14:1 ratio of G1 versus G2/M cells based on flow cytometry estimation, data not shown). To visualize cells in S-phase, cells were pulsed with 10 μM EdU for 30 mins before fixation. After fixation, a Click-It reaction of the EdU with Alexa488 azide was performed after the FISH hybridization and wash steps. The cumulative distributions for FISH signals from a high-slope region (yellow, right column) are shifted to the right compared to the internal control probe (magenta line) confirming the decondensed nature of DLCDs. In contrast, this shift was not seen for FISH probes targeting low-slope test regions. n≥ 300 alleles, 200 cells were imaged for each test probe.

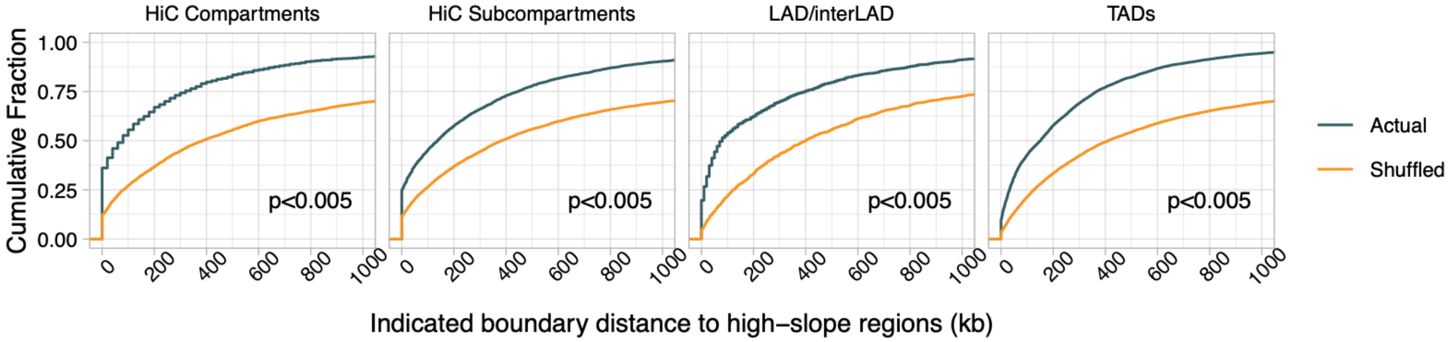


**Supplemental Figure S6 | DLCDs map near Chromatin Domain Boundaries**. Cumulative distribution showing fraction of Decondensed Large-scale Chromatin Domains (DLCDs) (y-axis) as function of distance in kb (x-axis) from (left to right): Hi-C compartments, Hi-C sub-compartments, Lamin B1 DamID LAD, and Hi-C TAD boundaries (green lines) as compared to shuffled boundaries (orange lines) in K562 cells. Boundary widths were 100 kbp for Hi-C compartments and sub-compartments, and LADs. P-values were measured empirically through shuffling DLCDs across the genome 200 times. Distances were measured from center of DLCDs (defined as regions in the top decile of TSA-seq slope values).

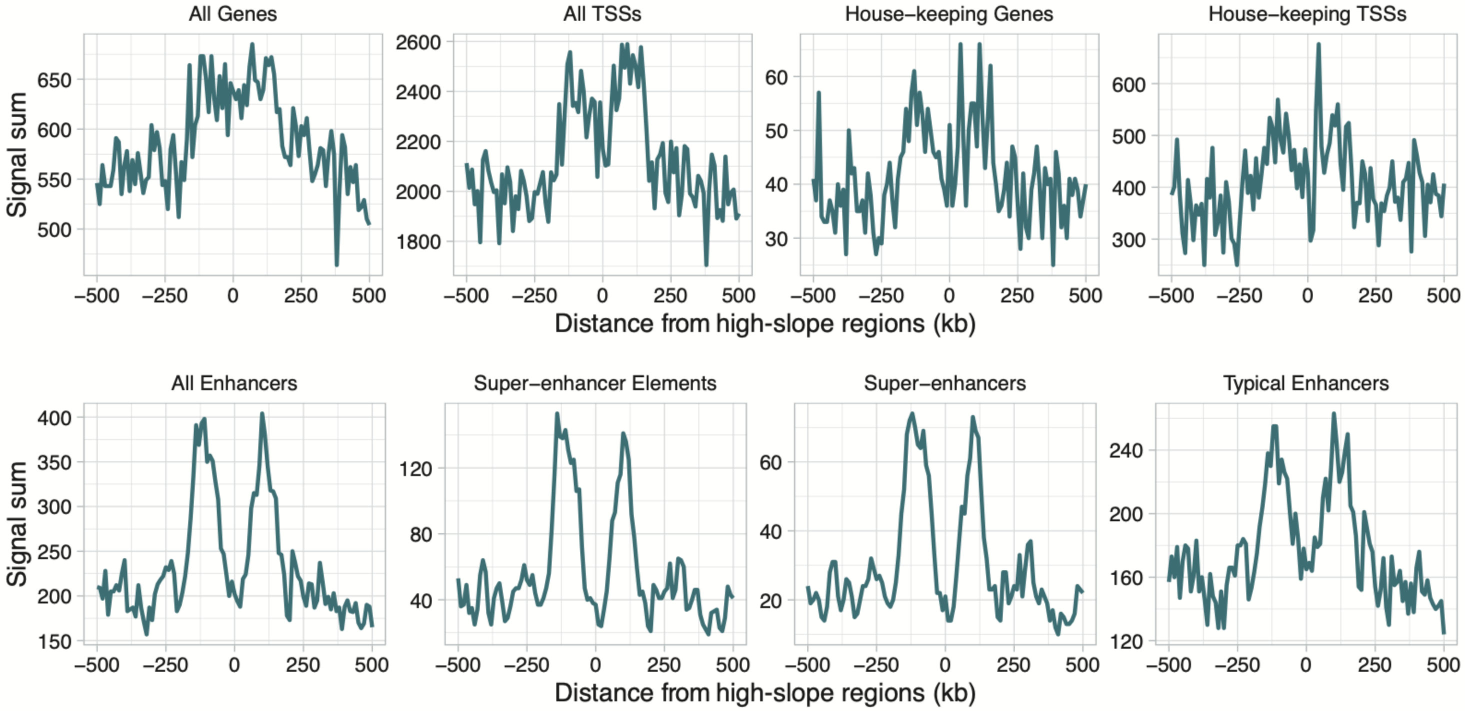


**Supplemental Figure S7 | DLCDs are flanked by super-enhancers, enhancers, transcription start sites (TSSs), and genes.** Profile plots of cis elements as a function of distance (+/- 500 kb, x-axis) from DLCD centers (0 kb). DLCDs were defined by the high-slope (top 10%) of the top decile of mean of TSA-seq slopes calculated across multiple TSA-seq datasets. Plots are shown (left to right, top to bottom) for all genes, all TSSs, house-keeping genes only, house-keeping TSSs, all enhancers, super-enhancer elements (defined as smaller enhancer), super-enhancers, and typical enhancers (enhancers that are not super-enhancers). The highest fold peak enrichment relative to background levels (~3-fold) are for super-enhancers and super-enhancer elements. Units (y-axis) are number of elements per 10 kb bin.

**Supplemental Figure S8 | DLCDs are flanked by transcription factors and chromatin associated proteins.** Profile plots of K562 ChIP-seq peak counts for various transcription and chromatin related proteins as a function of distance (+/- 500 kb, x-axis) from DLCD centers (0 kbp). DLCDs were defined by the high-slope (top 10%) of the top decile of mean of TSA-seq slopes calculated across multiple TSA-seq datasets. K562 DLCDs are centered at zero bp and the peak counts are summed within 10 kbp bins for +/-500 kbp flanking the DLCDs. Units (y-axis) are peak counts per bin.

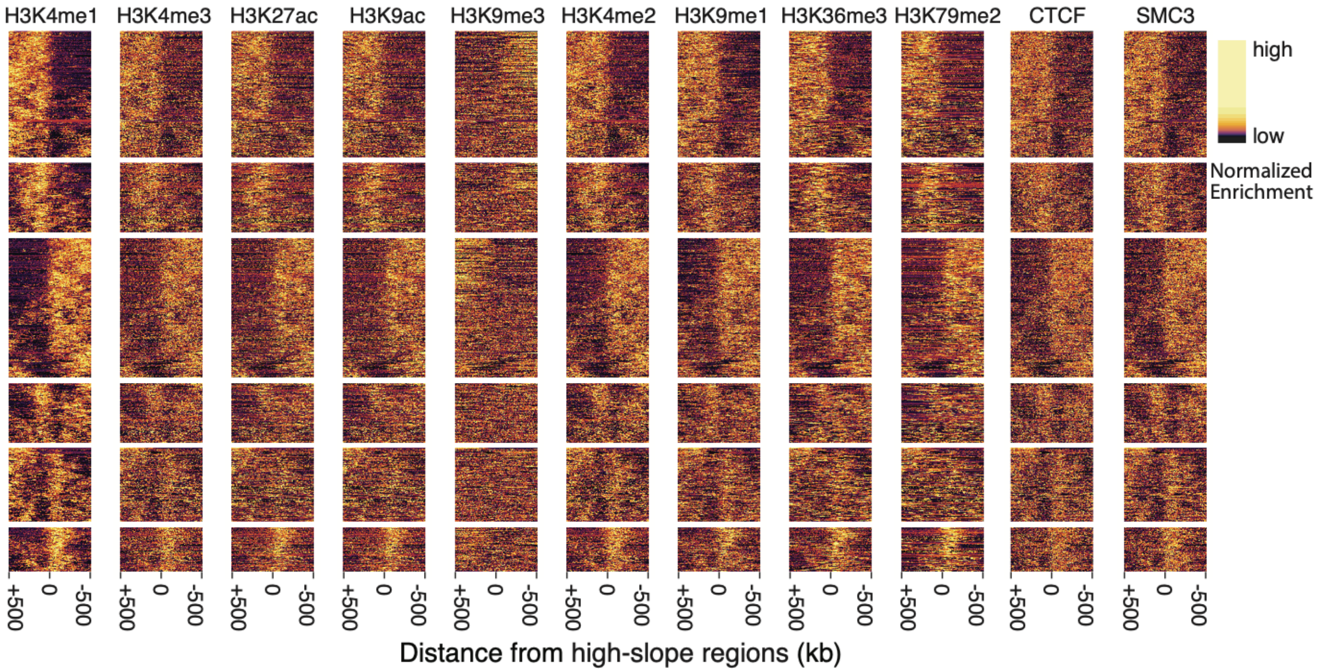


**Supplemental Figure S9 – Histone marks flanking individual DLCDs.** Heatmaps of ChIP-seq measurements of histone marks as function of distance (+/- 500 kbp) from DLCD centers. DLCDs (0 bp) were identified by the high-slope (top 10%) of the top decile of mean of TSA-seq slopes calculated across multiple TSA-seq datasets. Each heatmap shows the distribution of different histone marks in 10 kbp bins flanking DLCDs. ChIP-seq pulldown/input values across each row of each heatmap are normalized independently to emphasize the local variations in these marks. Rows in each heatmap are ordered according to a clustering scheme (see Methods) using H3K4me1 data.

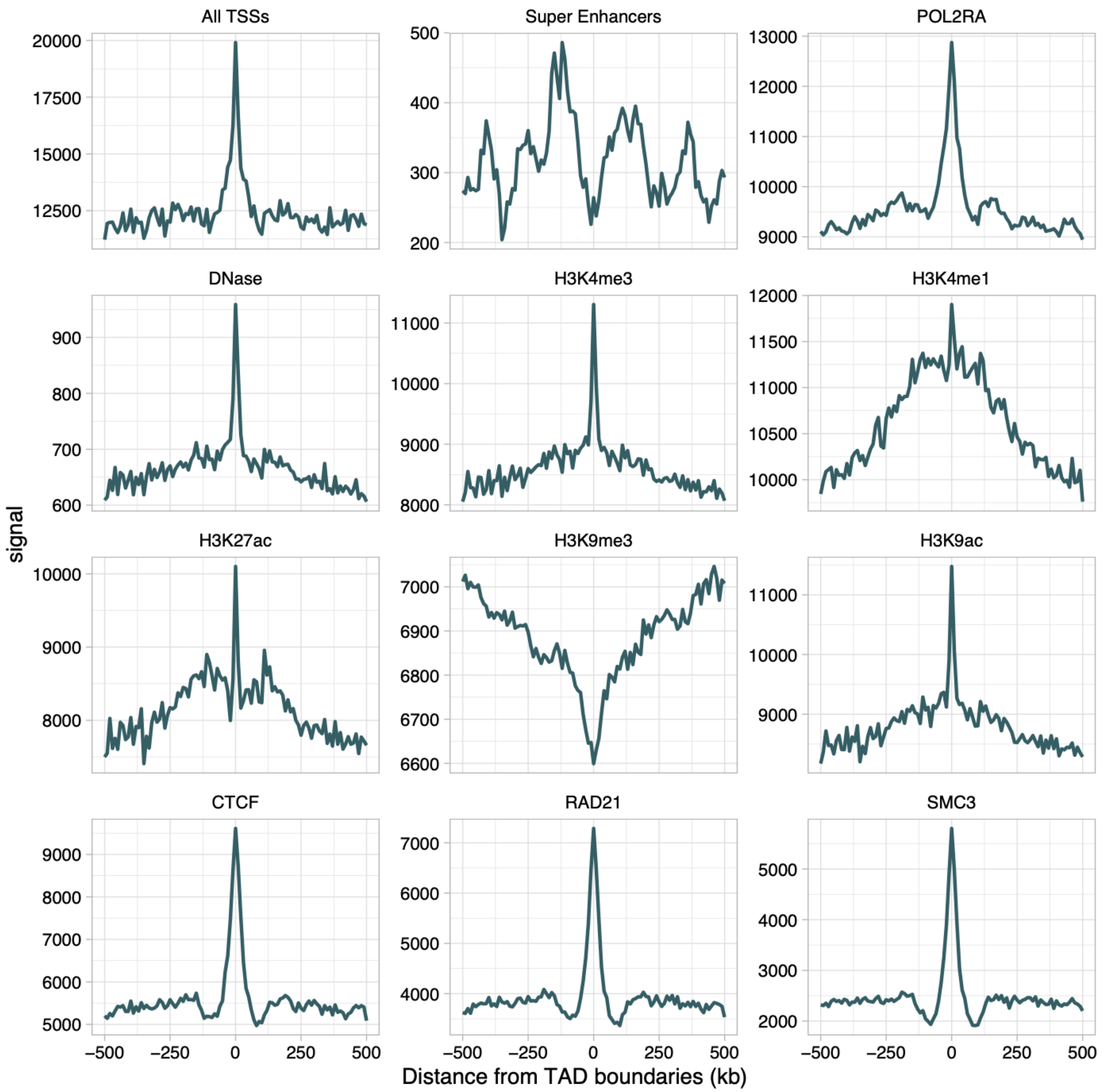


**Supplemental Figure S10 – Profile plots showing averaged levels of cis elements, chromatin marks, and genomic architectural proteins flanking TAD boundaries.** Profile plots in K562 cells of (left to right, top to bottom) All transcription start sites (TSSs), super-enhancers, POL2RA subunit of RNA pol 2, DNase 1 sensitivity, H3K4me3, H3K4me1, H3K27ac, H3K9me3, H3K9ac, CTCF, RAD21, SMC3 as function of distance (+/-500 kbp, x-axis) from TAD boundaries (0 bp). Data for TAD boundaries is from Rao et.al 2014. Units (y-axis) are peak counts per 10 kb bin for TSSs, super-enhancers, POLR2A, CTCF, RAD21, and SMC3 and pulldown/input ratios per bin for all other plots.

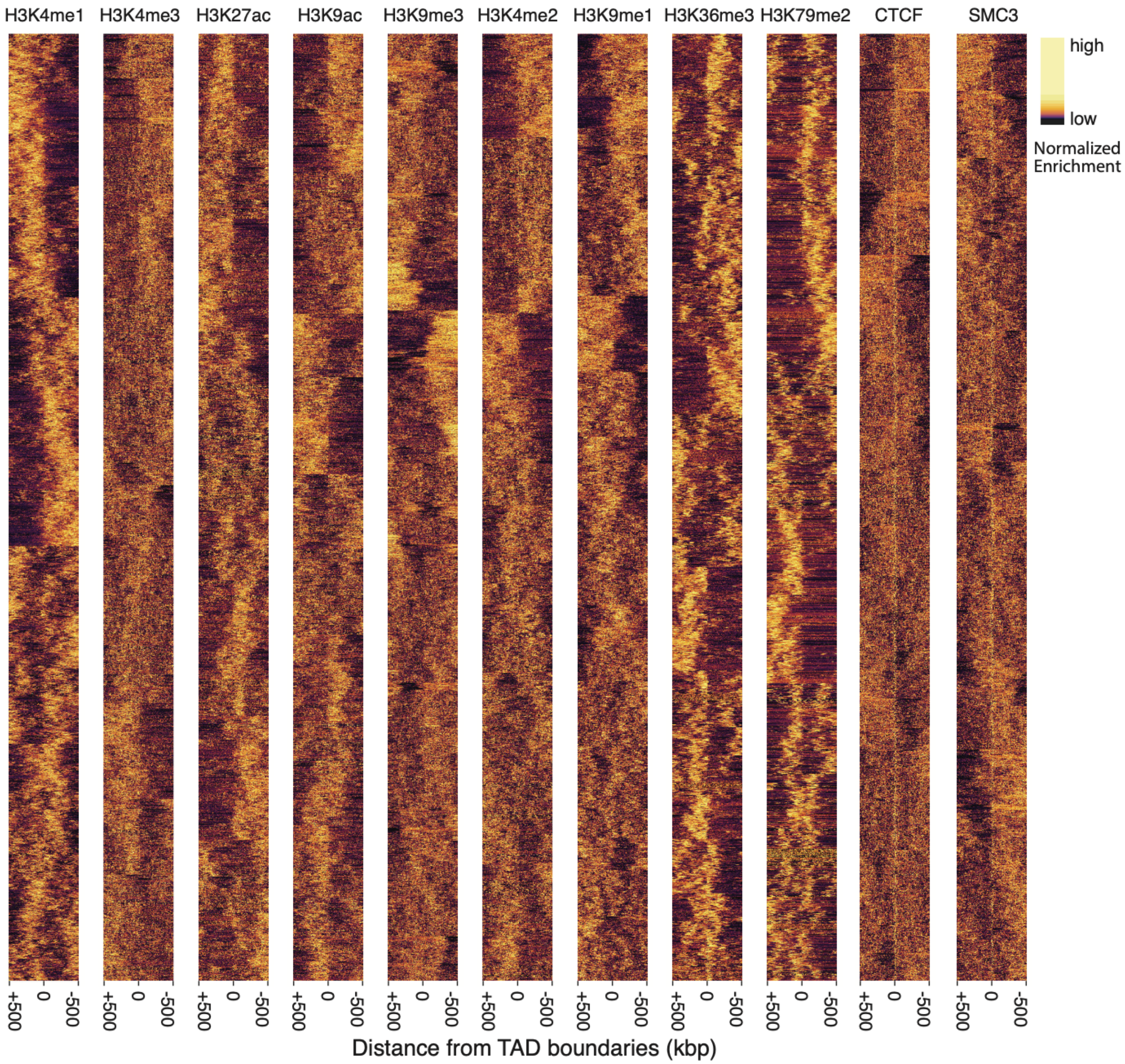


**Supplemental Figure S11 – Distribution of histone marks flanking individual TAD boundaries.** Heatmaps of histone marks (10 kbp bins) flanking (+/-500 kb, x-axis) TAD boundaries (0 bp) in K562 cells. Values across each row (pulldown/input) of each heatmap are normalized independently to emphasize the local variations in these marks. Rows in each heatmap are ordered according to a clustering scheme described in the method. This clustering was performed independently on each heatmap.
